# Moderately pathogenic maternal influenza A virus infection disrupts placental integrity but spares the fetal brain

**DOI:** 10.1101/2021.01.22.427849

**Authors:** Adrienne M. Antonson, Adam D. Kenney, Helen J. Chen, Kara N. Corps, Jacob S. Yount, Tamar L. Gur

**Author notes:** Co-corresponding author information: Adrienne M. Antonson, PhD, University of Illinois at Urbana-Champaign, 220 Edward R. Madigan Laboratory, 1201 W. Gregory Drive, Urbana, IL 61801 USA, Tamar L. Gur, MD PhD, The Ohio State University, 120A Institute for Behavioral Medicine Research Building, 460 Medical Center Drive, Columbus, OH 43210 USA. Declarations of interest: none.

## Abstract

Maternal infection during pregnancy is a known risk factor for offspring mental health disorders. Animal models of maternal immune activation (MIA) have implicated specific cellular and molecular etiologies of psychiatric illness, but most rely on pathogen mimetics. Here, we developed a mouse model of live H3N2 influenza A virus (IAV) infection during pregnancy that induces a robust inflammatory response but is sublethal to both dams and offspring. We observed lung inflammatory cytokine production and severely diminished weight gain in IAV-infected dams. This was accompanied by immune cell infiltration in the placenta and partial breakdown of placental integrity. However, indications of IL-17A signaling and fetal neuroinflammation, which are hallmarks of mimetic-induced MIA, were not detected. Our results suggest that mild or moderately pathogenic IAV infection during pregnancy does not inflame the developing fetal brain, and highlight the importance of live pathogen infection models for the study of MIA.

**Highlights:**

- A mouse model of influenza A virus (IAV) infection during pregnancy was established
- Moderate IAV infection induced lung inflammation and blunted weight gain in dams
- Maternal IAV infection caused mild pathology in the placenta without pup loss
- Moderate gestational IAV infection did not induce fetal brain inflammation
- An IAV infection severity threshold may exist for inducing fetal neuroinflammation

## 1.0 Introduction

Influenza A virus (IAV) infection during pregnancy poses a substantial risk for heightened morbidity and mortality in both mother and infant. Poor infant outcomes associated with maternal IAV infection, such as low birth weight, premature delivery, and stillbirth, occur in the absence of IAV vertical transmission, as the virus rarely breaches the maternal-fetal interface^1–4^. Given a lack of direct fetal infection, prolonged health complications due to prenatal exposure to maternal IAV infection, including development of neuropsychiatric disorders^5,6^, have mostly been attributed to the maternal inflammatory response. While there is evidence in humans that elevated maternal cytokines during pregnancy contribute to poor offspring psychiatric outcomes^7^, much of the data on prenatal inflammation comes from animal models of maternal immune activation (MIA)^8^. These models indicate that aberrant cytokine signaling can undermine placental integrity and disrupt homeostatic neurodevelopment prenatally^9–11^. However, it remains unclear whether there is a threshold of infection severity required for fetal neuroinflammation during MIA, particularly in IAV infections.

It is also important to note that the majority of MIA animal models utilize pathogen mimetics as immunostimulants, including bacterial endotoxin lipopolysaccharide (LPS) or synthetic viral dsRNA mimetic polyinosinic-polycytidylic acid (poly I:C). These non-replicating toll-like receptor (TLR) ligands trigger acute phase inflammatory responses that are short-lived (lasting 24-48 hours), and allow for the targeting of fetal developmental windows and precise control of cytokine responses through dose regulation^12,13^. While the enhanced control and reproducibility obtained with pathogen mimetics is advantageous^12^, the immune response is difficult to translate to that of a live pathogen infection. For example, inhalation of live influenza virus into the respiratory tract is followed by continuous viral replication and spread over multiple days, triggering a prolonged and robust inflammatory response involving multiple concurrently activated intracellular detection and signaling pathways^14^. Distinct inflammatory cascades, initiated by early innate responses^15^ and then later adaptive responses to IAV^16^, generally occur over the span of one to two weeks until the virus is cleared. Overall, immune responses to IAV infection during pregnancy extend across a more prolonged period and involve a wider array of cellular inflammatory and regulatory pathways as compared to MIA induced by a bolus of LPS or poly I:C. To date, comparatively few studies have been conducted using live IAV infection in the MIA field^17–19^.

While essential progress continues to be made in elucidating potential etiologies of disrupted fetal neurodevelopment during MIA, these mechanisms have yet to be confirmed in models that use live pathogen infections. In particular, recent evidence implicates maternal T helper 17 (T_H_17) cells and interleukin (IL)-17A as primary drivers of altered fetal brain development during poly I:C-induced MIA^9,20–22^. Genetic knock out of transcription factor retinoic acid receptor-related orphan nuclear receptor gamma t (RORγt), expressed by IL-17-producing cells and a master regulator of T_H_17 cell development^23^, can rescue MIA-induced cortical malformations^9^. Similarly, IL-17A blockade in pregnant dams protects the developing fetal brain from MIA-induced disruptions in morphology and proteostasis, and normalizes behavioral outcomes^9,22,24^. The majority of T_H_17 cells reside in the intestine, and endogenous segmented filamentous intestinal bacteria (SFB), which promote T_H_17 cell development^25^, have been shown to be required for the emergence of MIA-associated offspring phenotypes^20,22,24^. Notably, respiratory IAV infection also promotes T_H_17 cell development and IL-17A-dependent intestinal injury in reports studying non-pregnant male mice^26^. Furthermore, shifts in the intestinal microbiome are evident during respiratory IAV infection despite the fact that the virus does not infect intestinal tissue^27^. Disrupted microbial composition is concomitant with intestinal inflammation^28^ and altered production of antimicrobial peptides^29,30^, and microbe depletion prior to IAV inoculation has been shown to diminish IAV-induced IL-17A production and intestinal injury^26^. Thus, there is mounting evidence that IAV infection during pregnancy may elicit heightened levels of maternal IL-17A, similar to what is observed in poly I:C-induced MIA models, though this remains to be investigated.

Here, we sought to develop a maternal IAV infection mouse model using an H3N2 virus strain and dose that is well characterized to induce moderate pathogenicity in mice^31–34^. A model of moderate infection severity was desirable as this represents the majority of sublethal seasonal influenza virus infections in humans^35^. Our results show that, despite lung and systemic inflammatory responses to this IAV infection, fetal brain inflammation was not observed. Consistent with the prevalence of mild-to-moderate infections during pregnancy, these data demonstrate that not all infection-induced MIA results in fetal neuroinflammation, and instead supports a model wherein a high severity threshold must be reached to induce detrimental effects on fetal neurodevelopment. Our study emphasizes the need for live pathogen-induced MIA models to further parse out discrete mechanisms that have the potential to drive aberrant offspring outcomes.

## 2.0 Materials and Methods

### 2.1 Animals

Nulliparous adult C57BL/6NTac female mice were obtained from Taconic Biosciences (Cambridge City, IN), pair-housed, and allowed to acclimate to The Ohio State University Wexner Medical Center vivarium for one week. Two days prior to breeding, females were housed in the soiled cage of a male stud (to initiate estrous), and on the third day that male stud was added to the cage. Presence of a vaginal plug was designated as gestational day (GD) 1, and females were removed to clean cages after breeding. Animals were maintained under sterile conditions on a 12 h light-dark cycle and body weights were recorded daily. A total of 37 pregnant dams across four identical replicates were used in this study. All animal research was approved by and performed in accordance with The Institutional Animal Care and Use Committee at The Ohio State University.

### 2.2 Influenza A virus inoculation

Mouse-adapted influenza A virus strain X31 (H3N2) was propagated and titered in 10-day embryonated chicken eggs as previously described^31^. On GD10, pregnant dams were randomly assigned to IAV or Con groups and anesthetized by inhalation of isoflurane (Henry Schein Animal Health, Dublin, OH) before intranasal inoculation with IAV strain X31 at a dose of 1000 TCID_50_ sterile saline or mock inoculated with sterile saline. This dose was selected to mimic the minimal infectious dose of IAV in humans^36,37^, and we have previously validated mild-to-moderate infection severity of this same dose and strain in non-pregnant male mice^31^. A total of 20 pregnant dams were inoculated with X31 (IAV group) and 17 pregnant dams were mock-inoculated (Con group) across 4 identical replicates. The MIA Model Reporting Guidelines Checklist^19^ is included as **Supp. File A**.

### 2.3 Tissue collection

Seven days post inoculation (dpi; GD17), pregnant dams were euthanized by CO_2_ inhalation, and tissues were excised under sterile conditions. Whole blood (obtained through cardiac puncture) was collected into EDTA-coated tubes and stored on ice until centrifugation at 13,300 rpm at 4°C for 15 min; plasma was aliquoted and stored at −80°C until analysis. The uterus was removed and individual fetuses and placentas were dissected out; any fetal resorptions were noted. Two placentas per litter were preserved in 10% Neutral Buffered Formalin for histology. Maternal intestinal tissue was removed, cleaned of mesenteric fat, and colon length obtained. Cross sections of the colon and ileum were preserved in methacarn solution (vol/vol: 60% methanol, 30% chloroform, 10% acetic acid) for histology. The small intestine was flushed with sterile PBS, sliced longitudinally, and cut into 1 cm pieces stored in ice cold HBSS with 5% FBS until further processing. All other tissues were snap frozen and stored at −80°C until analysis.

### 2.4 Intestinal epithelial cell (IEC) and intraepithelial lymphocyte (IEL) selection

Sections of the small intestine were immediately transferred to a digestion solution (HBSS with 10mM HEPES, 5mM EDTA, 1mM DTT, 5% FBS) and incubated at 37°C for 20 min with slow rotation. Samples were then vortexed and supernatant (containing IECs and IELs) was passed through a 100 μm cell strainer and kept on ice. Intestinal sections were incubated and rotated in digestion solution two more times, vortexed, and supernatant passed through a 100 μm and then 70 μm cell strainer. The combined flow-through containing IECs and IELs was centrifuged at 400 × *g* for 10 min at 4°C, cells were washed in buffer (PBS with 2% FBS and 2 mM EDTA), passed through a 70 μm cell strainer, then centrifuged and resuspended in buffer containing CD45 MicroBeads (Miltenyi Biotec, Auburn, CA). Cells were incubated with microbeads for 15 min at 4°C, then washed with buffer and passed through LS columns for magnetic cell sorting, as per manufacturer’s protocol (Miltenyi Biotec). Both CD45 positive (IEL) and negative (IEC) cell fractions were collected and stored at −80°C until analysis.

### 2.5 Protein assays

Protein was isolated from tissues by sonication (40% amp) in lysis buffer (100:1 T-PER Tissue Protein Extraction Reagent and Halt Protease and Phosphatase Inhibitor Cocktail; Thermo Scientific, Waltham, MA). Homogenized samples were then incubated for 30 min at 4°C, centrifuged at 13,300 rpm at 4°C for 15 min, and protein lysates were aliquoted. Protein concentration was determined using the Pierce BCA Protein Assay kit (Thermo Scientific) and normalized prior to each assay.

Protein lysates from fetal brains (one per litter, normalized to 4 mg/mL) and placentas (one per litter, normalized to 7.4 mg/mL) were pooled to obtain a total of 1 mL per treatment group per tissue. Each pooled sample was assayed for 40 different cytokines using the Proteome Profiler Mouse Cytokine Array Panel A kit according to manufacturer’s instructions (R&D Systems, Minneapolis, MN). Membranes were imaged using a LI-COR Odyssey Fc chemiluminescence imager (Lincoln, NE) and mean pixel density between duplicate spots was quantified using ImageJ software (NIH, Bethesda, MD).

Protein lysates from lung tissue (normalized to 6 mg/mL using Diluent 41) and plasma samples (diluted 1:1 in Diluent 41) were assayed in duplicate on a custom multiplex assay (U-PLEX Assay Platform, Meso Scale Discovery, Rockville, MD) containing 9 cytokines: IFN-α, IFN-β, IFN-γ, IL-1β, IL-6, TNFα, IL-17A, IL-17C, IL-17E/IL-25. The assay was performed according to manufacturer’s instructions using Alternate Protocol 1, Extended Incubation. Concentration of circulating LPS-binding protein (LBP) was determined in plasma samples (diluted 1:5) using Mouse LBP ELISA Kit (Abcam, Cambridge, MA) according to manufacturer’s instructions. Protein lysates from colon tissue (normalized to 3.5 mg/mL) were assayed in duplicate using the Mouse IL-17 DuoSet ELISA (R&D Systems) according to manufacturer’s instructions.

### 2.6 Quantitative real-time PCR

RNA was isolated using TRIzol Reagent as per the TRI Reagent Protocol (Invitrogen, Carlsbad, CA). cDNA was synthesized using the High-Capacity cDNA Reverse Transcription Kit (Applied Biosystems, Foster City, CA), and PCR was performed on a QuantStudio 3 Real-Time PCR System machine using TaqMan Fast Advanced Master Mix (Applied Biosystems). Data were analyzed using the 2^−ΔΔCt^ method against housekeeping gene *Rpl19* and presented as relative expression compared to control, unless otherwise stated. Statistics were run on relative expression values. Primer information is listed in **Supp. Table S7**. Custom primers (Integrated DNA Technologies, Coralville, IA) against the IAV nucleoprotein (forward: 5’-CAGCCTAATCAGACCAAATG-3’; reverse: 5’-TACCTGCTTCTCAGTTCAAG-3’) were assayed using SYBR Green Master Mix (Applied Biosystems) to confirm viral presence in the maternal lung.

### 2.7 NanoString Gene Expression Panel

RNA was isolated from 12 fetal brains (one per litter) using TRIzol Reagent as per the TRI Reagent Protocol (Invitrogen), and delivered to the Genomics Shared Resource lab at The Ohio State University Comprehensive Cancer Center. RNA integrity was assessed using both BioAnalyzer and Qubit systems (mean RIN = 5.7), and the samples were assayed on the nCounter Neuroinflammation Panel (nanoString, Seattle, WA) according to manufacturer’s protocol. Raw data were analyzed using nSolver 4.0 Software (nanoString), applying the Advanced Analysis method with automated normalization gene selection. Automated probe exclusion using the “Omit Low Count Data” default selection resulted in omission of 166 probes. Based on quality control measures, two samples (one per treatment group) were identified as outliers and excluded from the analysis, resulting in 591 genes assayed across 10 samples. False discovery rate (FDR) correction for multiple comparisons was applied using Bonferroni correction.

### 2.8 Pathology

Histology services were performed in the Histology Laboratory of the Comparative Pathology and Digital Imaging Shared Resource (CPDISR) of The Ohio State University Comprehensive Cancer Center. Tissues were routinely processed for histopathology on a Leica Peloris 3 Tissue Processor (Leica Biosystems, Buffalo Grove, IL), embedded in paraffin, sectioned at an approximate thickness of 4-5 micrometers, and batch stained with hematoxylin and eosin (H&E) on a Leica ST5020 autostainer (Leica Biosystems) using a routine and quality controlled protocol. Slides were then evaluated by a board-certified veterinary comparative pathologist (Kara N. Corps, who was blinded to treatment groups) using a Nikon Eclipse Ci-L Upright Microscope (Nikon Instruments, Inc., Melville, NY). Representative photomicrographs were taken using an 18 megapixel Olympus SC180 microscope-mounted digital camera and cellSens imaging software (Olympus Life Science, Center Valley, PA).

Semi-quantitative scoring of placental histopathology findings was performed using a scoring rubric modified from previously published placental scoring methods^38,39^. Mineralization was scored using the following criteria: 0 = no mineral present beyond scattered small foci; 1 = rare or mild, multifocal mineralization; 2 = multifocal scattered foci of mineralization; 3 = large multifocal zones of mineralization or diffuse mineralization within a zone. Infiltration of neutrophils and mononuclear cells were scored using the following criteria: 0 = normal; 1 = slight increase; 2 = moderate increase; 3 = marked increase. Necrosis was scored using the following criteria: 0 = no necrosis present; 1 = multifocal necrosis exclusively in the maternal placenta; 2 = multifocal necrosis throughout the placenta; 3 = multifocal to coalescing, large zones of necrosis throughout the placenta; 4 = diffuse placental necrosis. Two sections each from two representative placentas per litter were scored and then averaged for each litter.

Semi-quantitative scoring of intestinal histopathology findings was performed using a scoring rubric adapted from established scoring methods^40,41^. Edema was scored using the following criteria: 0 = normal, 1 = mild/focal/single layer of ileum or colon, 2 = moderate/multifocal/multiple layers of ileum or colon, 3 = severe/widespread/transmural. Epithelial hyperplasia was scored using the following criteria: 0 = normal, 1 = slight/mild, 2 = moderate, 3 = marked: discrete nests of regenerated crypts delineated from adjacent mucosa with no obvious disruption of mucosal surface. Presence of neutrophils and lamina propria mononuclear cells were scored using the following criteria: 0 = normal, 1 = slight increase, 2 = moderate increase, 3 = marked increase. Crypt/gland inflammation was scored using the following criteria: 0 = normal mucosa, 1 = one-to-two inflammatory cells, 2 = cryptitis/significant gland inflammation affecting majority of structure, 3 = crypt/gland abscess/necrosis, 4 = crypt/gland absent due to mucosal destruction. Mucosal/crypt loss was scored using the following criteria: 0 = normal mucosa, 1 = shortening of basal one-third of crypts +/− slight inflammation and edema in the lamina propria, 2 = loss of basal two-thirds of crypts +/− moderate inflammation in the lamina propria, 3 = loss of all epithelium +/− severe inflammation in lamina propria or submucosa inflammation with surface epithelium intact/remaining, 4 = loss of all epithelium +/− severe inflammation in lamina propria +/− muscularis; exudate may cover damaged mucosa. Two sections of colon and ileum tissue per dam were scored and averaged.

### 2.9 Statistics

Dam (or litter) was treated as the experimental unit for all outcomes, and one representative fetus per litter was used for each experimental outcome, unless otherwise stated. All data (except nanoString gene expression data, described above) were analyzed using GraphPad Prism 8 software (San Diego, CA), with significance set at α = 0.05. Dam body weights across time were analyzed using repeated measures two-way ANOVA (time x infection), with Geisser-Greenhouse correction for sphericity and Bonferroni correction for multiple comparisons. Unpaired parametric t tests were used to compare Con and IAV groups, with Welch’s correction for unequal variances when applicable. For data that were not normally distributed (multiplex cytokine concentrations and histopathology scores), non-parametric Mann-Whitney tests were used. Outliers were identified and removed using the ROUT method with Q = 1%.

## 3.0 Results

### 3.1 Respiratory IAV infection during gestation induces a robust maternal inflammatory response

To determine whether the moderately pathogenic H3N2 IAV strain X31 infection could induce MIA, we inoculated pregnant dams intranasally with a dose of 1000 TCID_50_ on GD10. We have previously demonstrated that prenatal insults initiated at GD10 confer detrimental behavioral and neuroimmune outcomes in offspring^42–44^. Furthermore, murine infection beginning at GD10 approximates early gestation in the human, a time when the risk of IAV infection in determining aberrant offspring mental health outcomes is especially high^5^. We found that IAV infection severely blunted maternal body weight gain throughout the remainder of the gestational period, culminating in an average body mass that was 6.2 ± 1.2 g less than saline controls by GD17 (**Fig. 1A**). Indeed, 4 out of 20 IAV-infected dams even lost weight across the 7-day period (**Supp. Fig. S1A**). IAV infection did not affect litter size or number of fetal resorptions, demonstrating that blunted body weight gain was not due to fetal loss (**Supp. Table S1**). Further, GD17 maternal body weight was still stunted when normalized to litter size (**Supp. Fig. S1B**). When corrected for body weight, maternal spleen mass at 7 dpi (GD17) was increased in the IAV group, suggesting systemic expansion of immune cells in response to the virus (**Fig. 1B**). We confirmed the presence of viral RNA in infected maternal lungs through quantitative PCR (**Fig. 1C**). Within the lungs, considerable increases in secreted pro-inflammatory cytokines in response to IAV infection were detected, including classical inflammatory mediators IL-1β, IL-6, and TNFα, IFN-α, and IFN-γ, as well as T_H_17 cytokines, IL-17A and IL-17C (**Fig. 1D**; IFN-β and IL-17E were below detectable levels). Circulating levels of plasma IL-6, TNFα, and IFN-γ were also increased in response to IAV infection (**Fig. 1E**), but all other cytokines (IFN-α, IFN-β, IL-1β, and IL-17s) were outside the detectable range. Overall, these results demonstrate that X31 IAV infection of pregnant dams induces a significant inflammatory response and prevents healthy maternal weight gain without loss of pups, establishing this infection system as a moderate MIA model.

**Figure 1.**
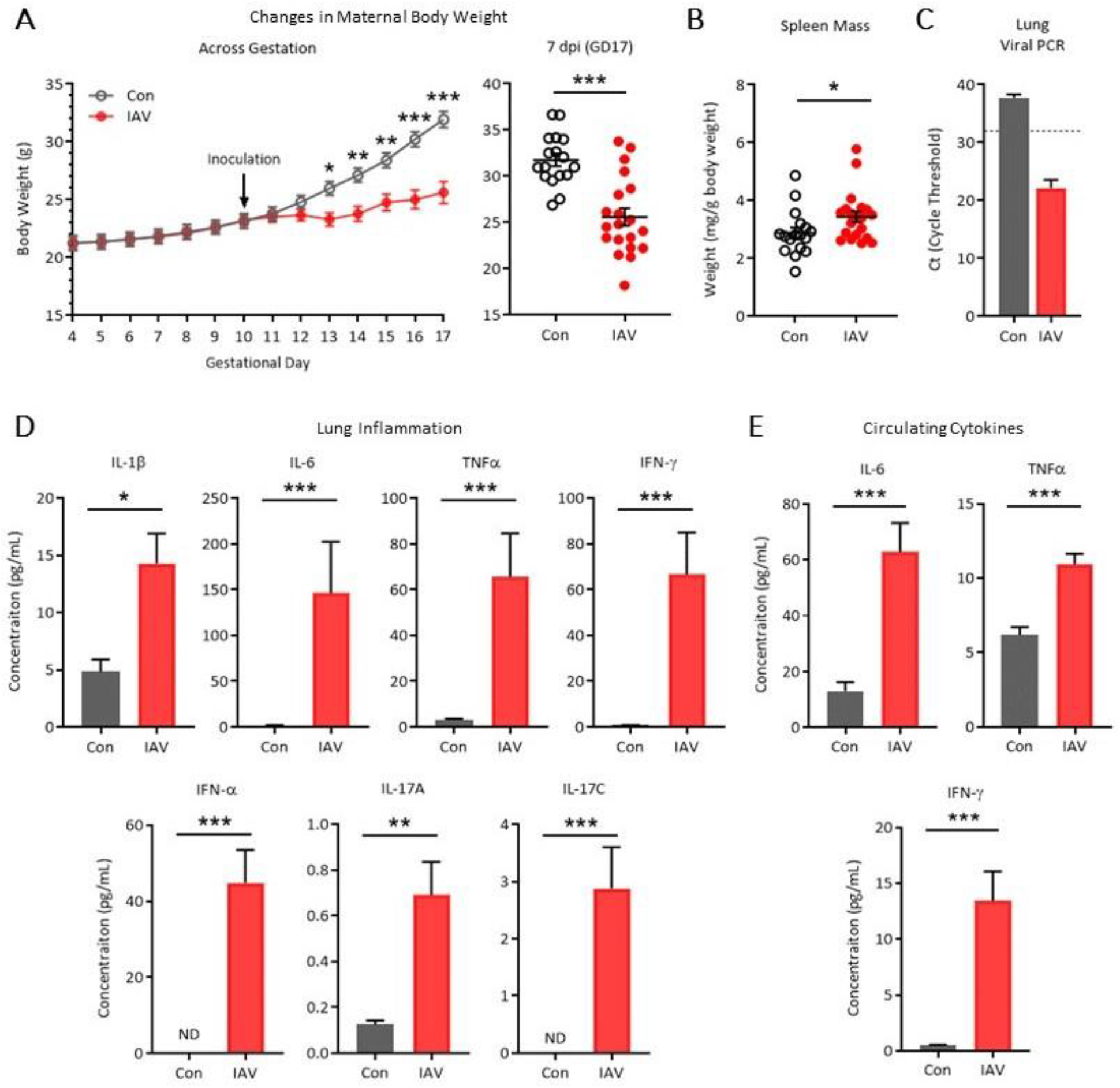
Gestational respiratory IAV infection induces robust systemic inflammation. IAV inoculation at gestational day (GD)10 **(A)** suppressed body weight gain beginning three days post-inoculation (repeated measures ANOVA: time x infection, p < 0.001, Bonferroni multiple comparisons correction, *** = adj p < 0.001, ** = adj p < 0.01, * = adj p < 0.05), resulting in substantial reduction in maternal body weight compared to control dams at 7 dpi. **(B)** Relative to body weight, maternal spleen mass at 7 dpi was increased due to IAV infection. Con: n = 17, IAV: n = 20. **(C)** Presence of IAV in the lung was confirmed through PCR, using a cycle threshold of ≤ 32 cycles as a confirmed infection. **(D)** Inflammatory cytokines associated with innate and adaptive anti-viral immune response were increased in the lungs of IAV infected dams at 7 dpi, concomitant with **(E)** increases in circulating IL-6, TNFα, and IFN-γ. Lung cytokines: Con n = 8, IAV n = 6; plasma cytokines: n = 13/group. IAV = influenza A virus; dpi = days post-inoculation; ND = non-detectable; *** = p < 0.001, ** = p < 0.01, * = p < 0.05.

### 3.2 Inflammation and antimicrobial responses extend to the placenta during maternal IAV infection

Given prior reports that placental disruptions can be induced by MIA^4,45^, we next examined whether maternal respiratory IAV X31 infection had downstream effects on the placenta. We scored H&E-stained placental sections for inflammatory lesions, and found increased incidence of necrosis, multifocal neutrophil infiltration, and mild-to-moderate mineralization in placentas from IAV-infected dams that was evident in both maternal and fetal layers (**Fig. 2A-F**). In order to broadly characterize the molecular immune profile of the placenta, we pooled samples of normalized protein extracts from placentas (one per litter) of control or IAV dams to obtain sufficient material to utilize a Proteome Profiler Cytokine Array Panel, which assessed 40 cytokines. Placentas from IAV infected dams showed modest increases in complement component C5/C5a, IL-1α, IL-1ra, and neutrophil chemoattractant KC/CXCL1 (**Fig. 3A**). Roughly 2-fold increases were observed for monocyte chemoattractant CCL2, T cell chemoattractant and CD4 ligand IL-16, and tissue inhibitor of metalloproteinases 1 (TIMP-1; **Fig. 3A**), consistent with the mild-to-moderate histological changes reported in **Fig. 2**. We next probed a select set of 26 genes of interest to examine transcription of classical inflammatory mediators implicated in MIA, as well as anti-viral and anti-bacterial response genes, and genes involved in placental barrier integrity (presented in **Supp. Table S2**). Of these, relative expression of three genes increased 2-fold (*Ifng*, bactericidal lectin *Reg3b*, and cytotoxic *Gzmb*; **Fig. 3B**), indicating a modest enhancement of antimicrobial responses. Expression of the remaining immune and barrier integrity genes were comparatively unchanged by maternal IAV infection (**Supp. Table S2**). Previous reports have suggested that type I IFN signaling during pregnancy can lead to disruptions of placental structural formation^46–48^. Thus, we also included interferon-responsive genes, *Ifitm3* and *Irf7*, but did not observe their placental upregulation upon IAV infection (**Supp. Table S2**), suggesting that the observed placental changes may not be driven by type I IFN signaling. Notably, relative expression of a subset of these same immune and antimicrobial genes was also largely unchanged in the uterus (**Supp. Table S3**). Together, our data suggest that maternal X31 infection leads to downstream immune and antimicrobial signaling and mild-to-moderate tissue breakdown in the placenta.

**Figure 2.**
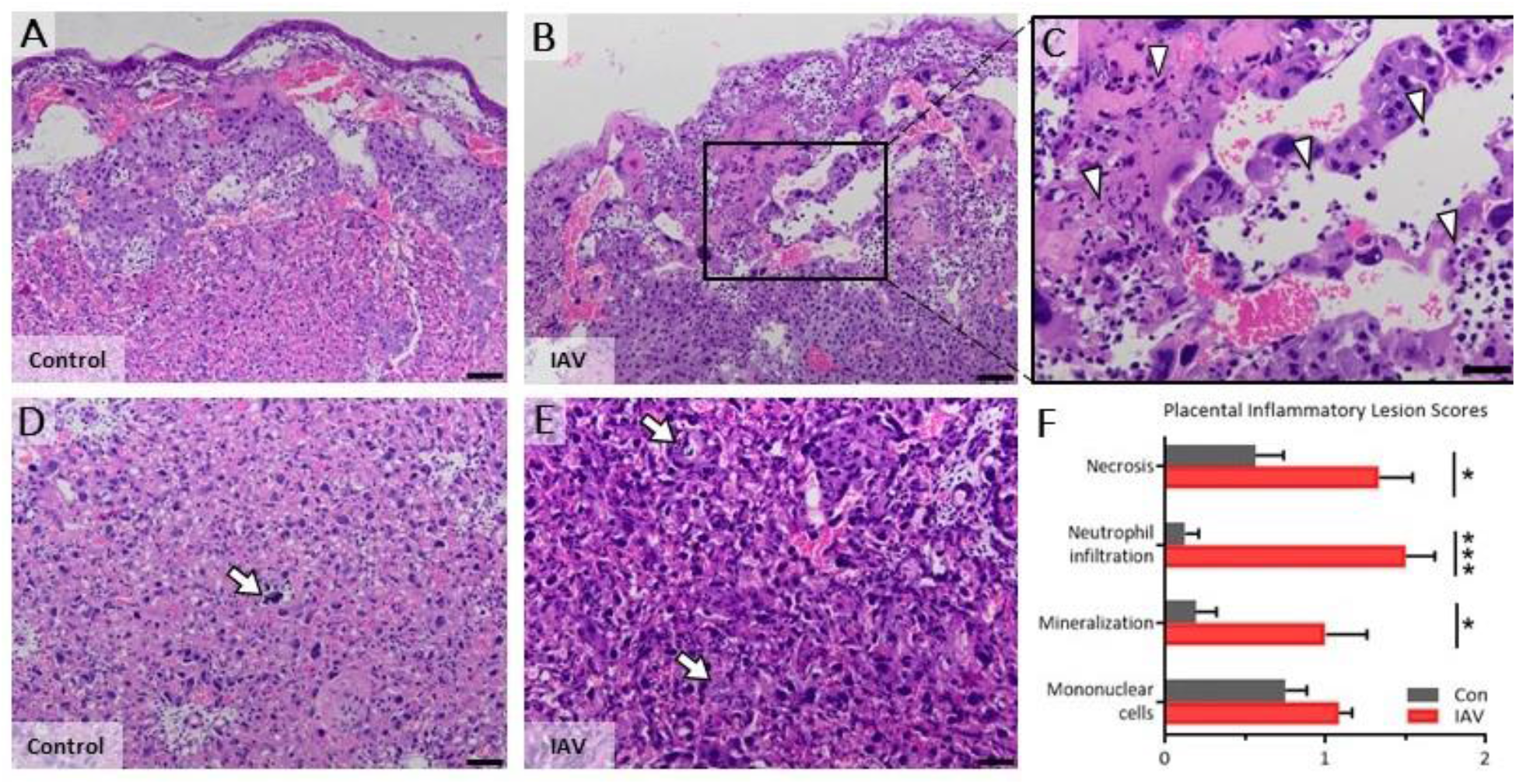
Maternal respiratory IAV-infection increases placental necrosis, mineralization and inflammation. Representative photomicrographs of H&E-stained placental sections and lesion score quantification from Control and IAV-infected dams. While Con dams had small foci of necrosis only in the maternal portion of the placenta and lacked an overt inflammatory response **(A)**, all IAV-infected dams had larger, multifocal to coalescing areas of necrosis in the maternal layers **(B)** and 2 of 6 had necrosis in the fetal layers. Necrosis in IAV dams was frequently accompanied by neutrophilic inflammation, and scattered neutrophils were present in all layers of the maternal tissue (**C**, arrows). Control dams had small foci of mineralization in the fetal vascular labyrinth, a frequent and expected finding (**D**, arrow). In contrast, IAV-infected dams had numerous scattered foci of mineral throughout the fetal vascular labyrinth that were occasionally accompanied by neutrophilic inflammation or focal necrosis (**E**, arrows). Increased overall basophilic staining was also evident **(B, E)** and is caused by a combination of increased circulating inflammatory cells, particularly mononuclear cells and multifocal neutrophils, and mild-to-moderate mineralization of capillary walls. Inflammatory lesion scores are quantified in **(F)**; scoring criteria is listed in the Methods. Two sections each from two representative placentas per litter were scored and averaged; Con: n = 8 litters, IAV: n = 6 litters. **A-B**: 10X magnification, scale bar = 100 μm; **C-E**: 20X magnification, scale bar = 50 μm. IAV = influenza A virus; *** = p < 0.001, * = p < 0.05.

**Figure 3.**
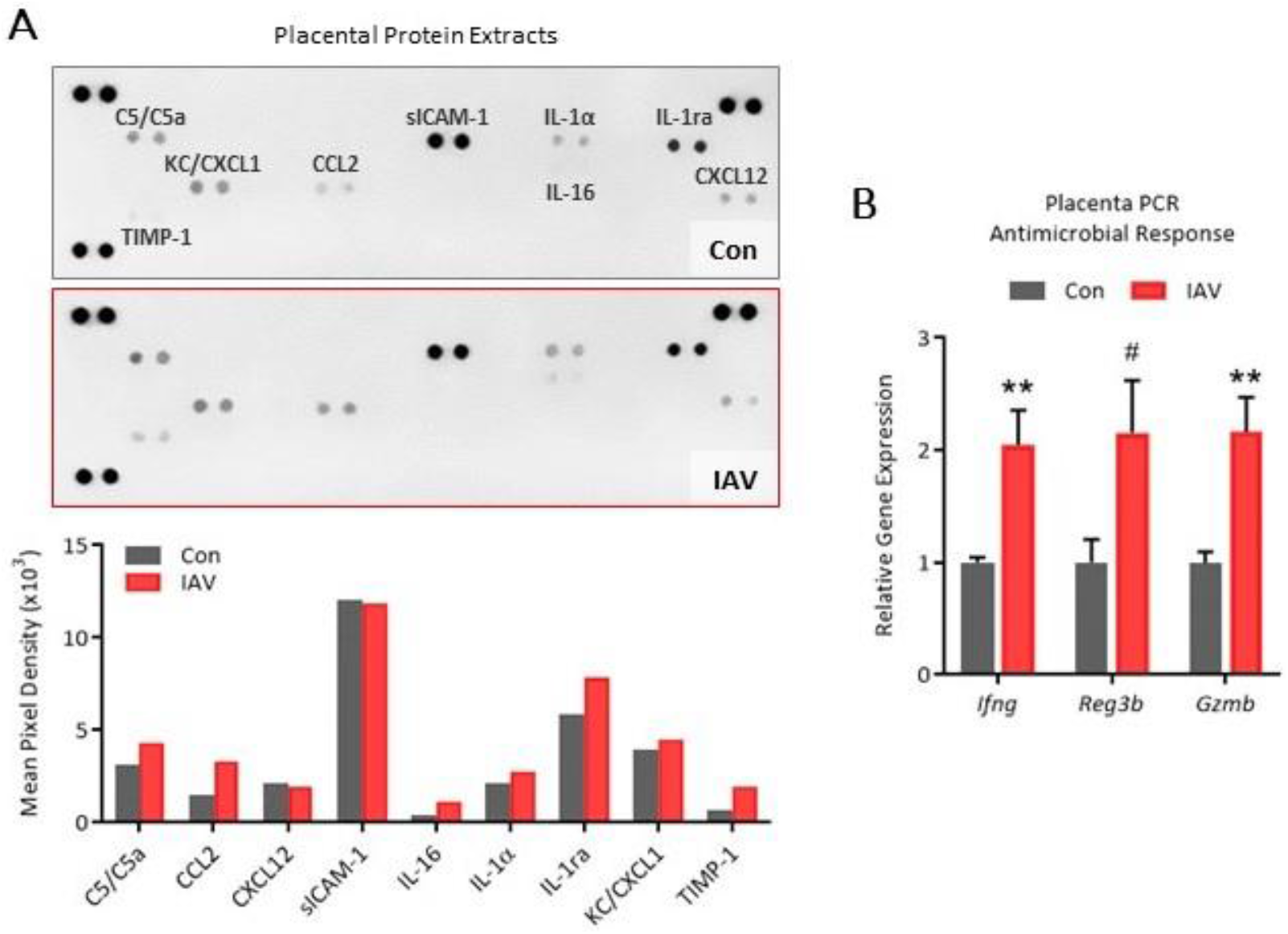
Maternal IAV infection increases placental inflammatory signaling and bactericidal responses. **(A)** Pooled placental protein extracts from control (n = 8) and IAV-infected (n = 6) dams were assayed on the Proteome Profiler Cytokine Array, suggesting enhanced immune responses due to maternal IAV infection. **(B)** Upregulation of *Ifng*, bactericidal C-type lectin *Reg3b*, and *Gzmb* (Granzyme B) expression in placentas from IAV-infected dams (n = 9-13/group). IAV = influenza A virus; ** = p < 0.01, # = p ≤ 0.10.

### 3.3 Fetal neuroinflammation is not overtly apparent during maternal IAV infection

Given that we observed a partial breakdown of placental integrity during maternal IAV infection, we further investigated whether immune homeostasis in the fetal brain was also affected. Cytokine array profiling of pooled fetal brain protein extracts revealed only two secreted proteins within detectable range, lymphocyte chemoattractant CXCL12 and soluble intercellular adhesion molecule-1 (sICAM-1), and these were at similar levels between groups (**Fig. 4A**). To more thoroughly examine the impacts of maternal IAV infection on neuroinflammation of the developing fetal brain, we investigated an extensive set of genes of interest across ten fetal brain samples using the nanoString nCounter Mouse Neuroinflammation Panel (a heatmap of the normalized data is presented in **Supp. Fig. S2**; log2 fold change and statistical results are presented in **Supp. Table S4**). However, out of 591 genes, we identified none that were differentially expressed in the fetal brain due to maternal IAV infection. To follow up on these findings, we conducted independent qPCR experiments to investigate relative expression of 23 genes implicated during prenatal insults and involved in neuroimmune responses, blood brain barrier integrity, and neurodevelopment (**Fig. 4B** and **Supp. Table S5**). We confirmed an overall lack of gene expression changes in the fetal brain upon maternal IAV infection, with only a modest but significant downregulation in *Il17ra* (encoding IL-17 Receptor A) and *Il1b*; *Il17a* failed to amplify within the fetal brain. In all, the notable absence of fetal neuroinflammation during moderately severe maternal IAV infection suggests that an infection severity threshold must be reached in order to induce immune imbalance in the developing fetal brain.

**Figure 4.**
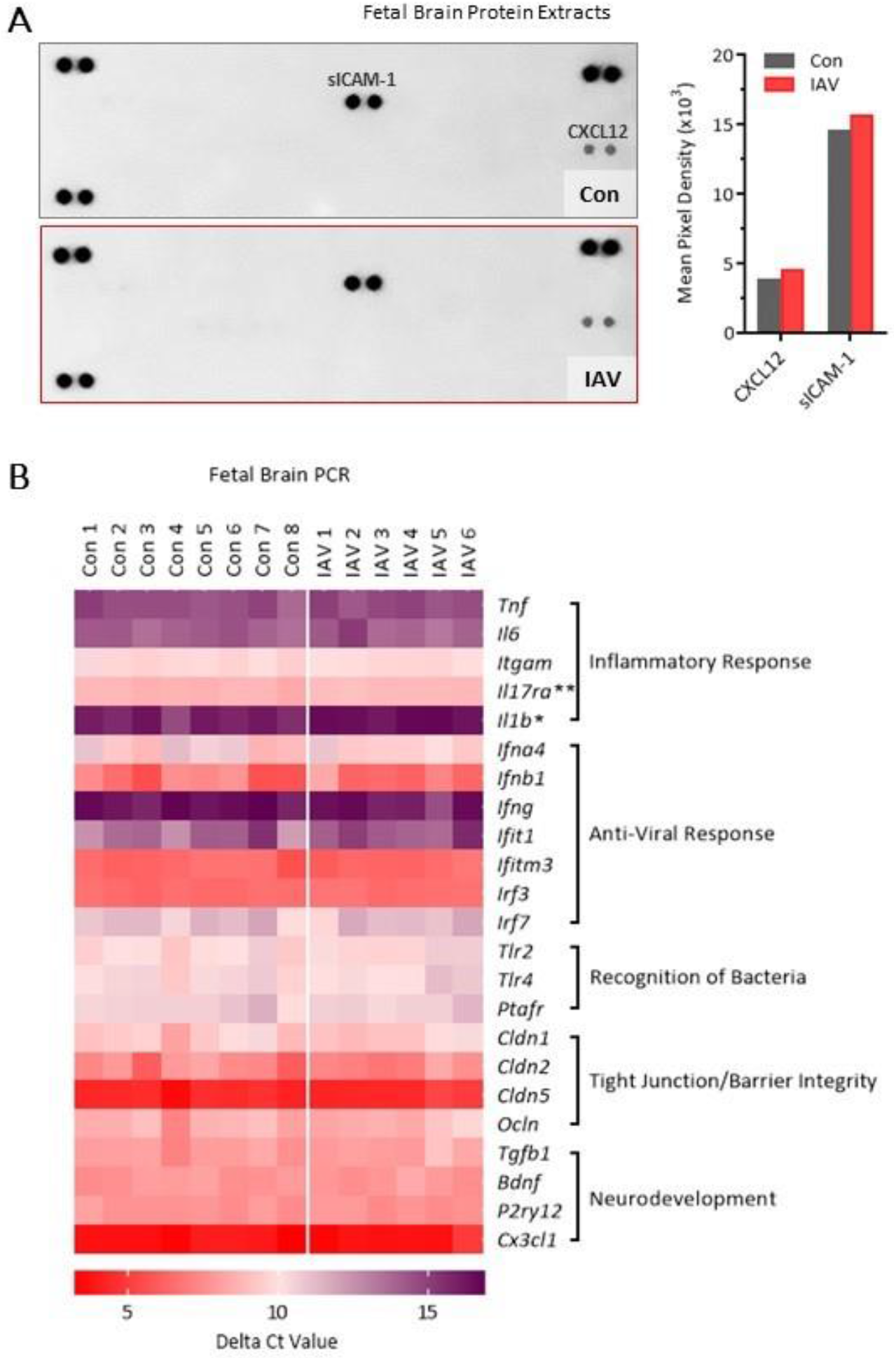
Fetal neuroinflammation is not overtly apparent during mild maternal IAV infection. **(A)** Pooled fetal brain protein extracts from litters of control (n = 7) and IAV-infected (n = 4) dams assayed on the Proteome Profiler Cytokine Array revealed only two detectable signaling molecules, suggesting limited inflammatory response to maternal IAV infection in the fetal brain. **(B)** Independent qPCR was carried out on fetal brain tissue to confirm and extend upon the nanoString Neuroinflammation Panel data. PCR data are represented as a heatmap of delta Ct values (compared to housekeeping control gene = *Rpl19*) of select genes involved in neuroimmune responses, blood brain barrier integrity, and neurodevelopment within the fetal brain. Purple indicates low expression compared to *Rpl19* (i.e. higher cycle threshold); red indicates high expression compared to *Rpl19* (i.e. lower cycle threshold). A modest but significant downregulation in *Il17ra* and *Il1b* expression was observed due to maternal IAV-infection, but no other genes were differentially expressed. Mean ± SEM and p-values for all genes are listed in Supp. Table S5. IAV = influenza A virus. ** = p < 0.01, * = p < 0.05.

### 3.4 Respiratory IAV infection has downstream consequences for maternal intestinal health

Since maternal IL-17A signaling has been associated with neurodevelopmental and behavioral abnormalities in offspring^9, 20, 22^, we sought to investigate whether downstream maternal IL-17A signaling is evident during IAV infection. We thus examined the maternal intestine for indications of intestinal immune injury and T_H_17 cell activation. As colon shortening is a hallmark indication of colitis^49^, we first measured colon length, which was shortened in IAV-infected pregnant dams compared to control (Fig. 5A). Relative expression of colonic *Ifng* and *Rorc* (which encodes T-cell dependent transcription factor RORγ) was upregulated by respiratory IAV infection, concurrent with a tendency for upregulated tight junction protein transcripts *Cldn1* and *Ocln* (p < 0.10), but not *Cldn2* or *Cldn5* (**Fig. 5A**). IL-17 protein within colon protein extracts did not differ (**Fig. 5A**). In order to follow up on the potential incidence of colitis, sections of colon and small intestine (ileum) tissue were H&E stained and examined for inflammatory lesions. However, we observed no differences in intestinal pathology between control and IAV-infected dams, with almost all mean lesion scores falling at or below 1 (i.e. normal-to-mild pathology; **Supp. Fig. S3**). We then isolated the CD45^+^ cell fraction (intraepithelial lymphocytes, IELs) of the small intestine to examine T_H_17-related genes. Again, while *Rorc* expression was increased in IELs of IAV-infected dams (**Fig. 5B**), there were no significant changes in expression of *Il17a* or *Il17ra*, nor were other T cell-related genes impacted, i.e. interferons (*Ifna4*, *Ifnb1*, *Ifng*) and *Il6* (**Supp. Table S6**). As T_H_17 cell differentiation is induced by shifts in commensal microbe populations^26,50^, and intestinal epithial cells (IECs, CD45^−^ fraction) mediate immune responses to endogenous microbes, we examined antimicrobial gene transcription within IECs as an indication of microbial dysbiosis. Expression of antimicrobial peptides alpha defensin 1 (*Defa1*), C-type lectins *Reg3b* and *Reg3g*, and dual oxidase *Duoxa2* were differentially impacted by IAV infection (**Fig. 5C**), indicating a disruption in IEC-controlled bacterial segregation. IEC expression of toll-like receptor 4 (*Tlr4*) and LPS-binding protein (*Lbp*), which detect bacterial LPS and help regulate antimicrobial molecule production, were downregulated while TLR4 signal transducer *Myd88* was unchanged (**Fig. 5C**). Relative expression of *Il17ra*, which binds IL-17A from T_H_17 cells and also aids in antimicrobial molecule production, was unchanged, as were *Tgfb1* and *Il23a*, which promote T_H_17 cell polarization (**Fig. 5C**). As a proxy for intestinal breakdown leading to release of bacteria into circulation, we measured LPS-binding protein in maternal plasma but found no differences between groups (**Fig. 5D**), consistent with maintained or enhanced tight junction protein transcripts within the colon (**Fig. 5A**). Overall, respiratory IAV X31 infection appears to drive mild downstream intestinal immune and microbial dysregulation, and while we observed consistently increased expression of transcription factor RORγ, indications of augmented IL-17 signaling were markedly absent. Given the known role of IL-17A in development of fetal neuroinflammation in MIA, the lack of intestinal IL-17 signaling in our IAV X31 infection experiments is consistent with the absence of fetal brain inflammation.

**Figure 5.**
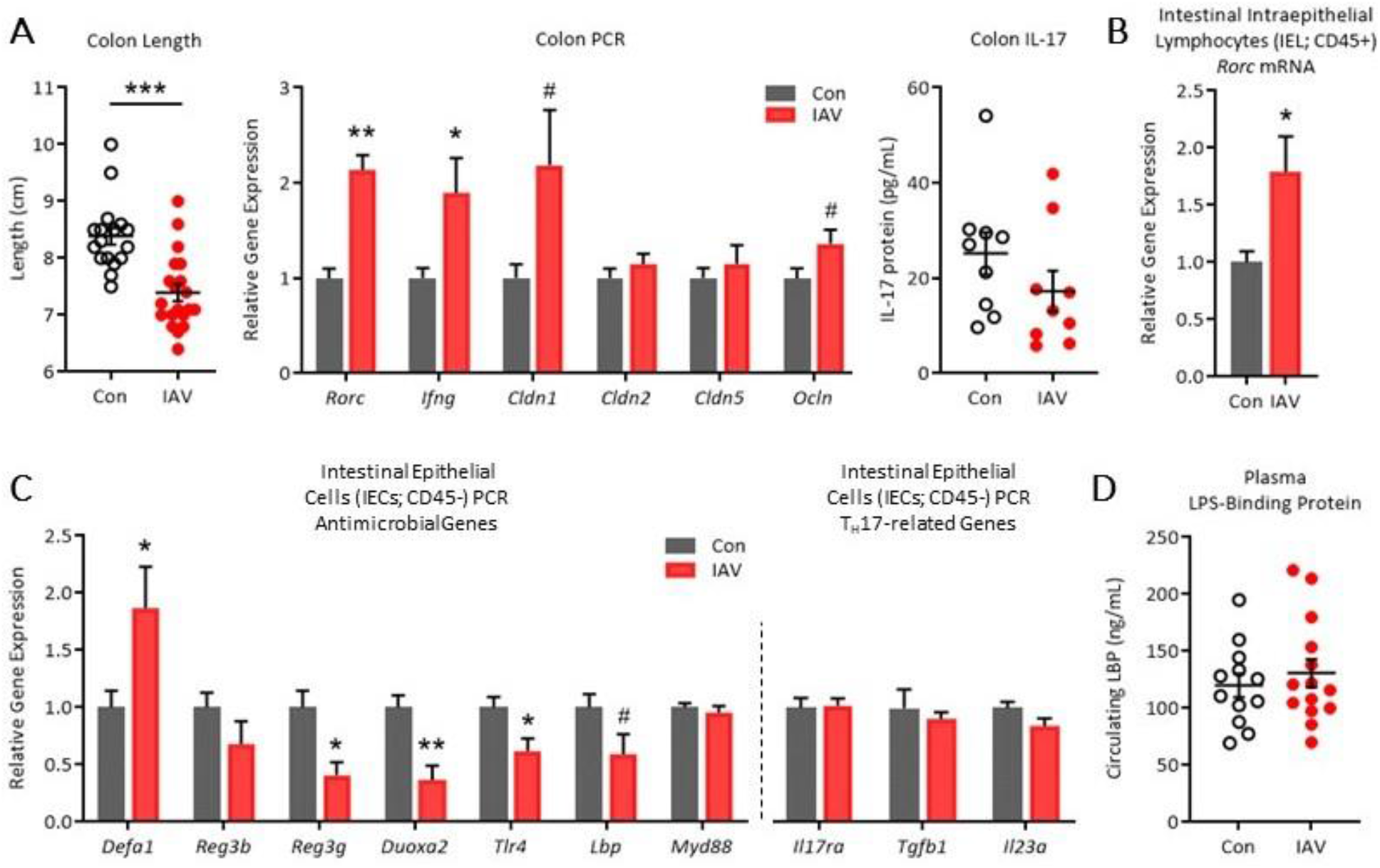
Downstream inflammatory impacts of gestational IAV infection on the maternal intestine. IAV-infected dams displayed **(A)** reduced colon lengths (Con: n = 17, IAV: n = 20) concurrent with upregulation in colonic expression of transcription factor *Rorc*, *Ifng*, and tight junction proteins *Cldn1* and *Ocln* (n = 6-13/group), while there were no changes in *Cldn2* and *Cldn5* expression or IL-17 protein within colon protein extracts (n = 9/group). **(B)** Intraepithelial lymphocytes (IEL; CD45+ fraction) isolated from the small intestine upregulated *Rorc* expression (n = 5-8/group). **(C)** Relative expression of most antimicrobial response genes was dysregulated in intestinal epithelial cells (IEC; CD45-fraction) of IAV-infected dam small intestine, while TH17-related genes were unchanged (n = 5-8/group). **(D)** The plasma concentration of LPS-binding protein, a proxy for disrupted intestinal barrier integrity, was not impacted by IAV infection (p = 0.52; Con: n = 12, IAV: n = 14). IAV = influenza A virus; *** = p < 0.001, ** = p < 0.01, * = p < 0.05, # = p ≤ 0.10.

## 4.0 Discussion

To address a major limitation of mimetic-induced MIA—the inability to capture a full pathogen-induced spectrum of inflammatory and regulatory immune responses—we developed a clinically relevant IAV infection model to probe the sequelae of MIA. Utilization of a moderately pathogenic IAV strain and dose, which is pertinent to the human condition, helps advance our understanding of the mechanisms mediating adverse fetal outcomes during seasonal IAV infection. Our data reveal that moderately severe IAV infection leads to maternal inflammation that is not restricted to the respiratory tract but also elicits off-target effects on the intestine and placenta. The notable absence of IL-17 signaling and of overt fetal brain inflammation supports the theory that an infection severity threshold likely exists for some MIA-mediated outcomes.

The clinically relevant moderate dose of X31 we selected is approximately 10 times lower than comparable studies [44], yet our IAV infection model still blunted maternal weight gain without impacting litter size. Others have shown that maternal IAV infection can slow placental and fetal growth, but does not increase fetal resorption^32,45,51^, in agreement with what we report here. While we did not record placental and fetal weights, it is likely that a similar growth retardation is present in our model considering the substantial body weight reduction in pregnant dams after correcting for litter size. Indeed, rates of intrauterine growth restriction leading to lower birth weight infants are increased following gestational IAV infection in humans^52^.

Although viral transmission to placental tissue during murine IAV infection is rare^3,4,45,51^, we observed mild-to-moderate placental pathology within all IAV litters, similar to a previous report^4^. Infiltration of neutrophils, but not mononuclear cells, was especially apparent and is common to both bacterial-induced intrauterine inflammation^53,54^ and preeclampsia^55^. Recent evidence suggests that maternal IAV infection can result in a vascular signature that mimics preeclampsia, albeit this study used a much higher dose of X31^45^. In our model, changes in placental transcription of matrix metalloproteinases 2 and 9, implicated in preeclampsia^56^, did not reach significance, although augmented secretion of metalloproteinase inhibitor TIMP-1^57^ was detectable in placental lysates. Additionally, increased placental mineralization, as observed here, can be an indication of placental vascular malperfusion^58^. Thus, it is possible that a mild preeclampsia-like phenotype also exists in our maternal IAV infection model, but comprehensive measurements of vascular alterations are still required.

On the other hand, upregulation of antimicrobial transcripts within the placenta also suggest that neutrophil trafficking might correspond to infiltration of endogenous bacteria, although the concept of *in utero* bacterial colonization is extremely controversial^59,60^. While vertical transmission of bacteria or their products during pregnancy has been demonstrated^61,62^, the notion that maternal IAV infection may lead to bacterial translocation to the placenta requires extensive follow-up. Interestingly, while there is increased necrotic tissue breakdown in placentas of IAV litters, barrier integrity may still be intact at 7 dpi considering that transcripts of tight junction proteins were unchanged. Further indication of a relatively resilient placental barrier system may be inferred from the lack of inflammatory signaling in the fetal brain, despite significant maternal inflammation in the lung and systemically. Indeed, recent evidence has identified nonclassical estrogen receptor GPER1, which is constitutively expressed in all major zones of the placenta, as a primary regulator of IFN signaling that protects fetal health during maternal IAV infection^51^. The lack of placental upregulation of interferon-response genes in our model indicates that GPER1 activation is likely suppressing maternally-derived IFN signaling within fetal tissues. Outside of IFNs, previous work has demonstrated the essential involvement of cytokine IL-6 in evoking neurodevelopmental abnormalities during mimetic-induced MIA^63–65^, which revealed the necessity of transplacental IL-6 signaling^11^. While IL-6 was increased approximately 3-fold in maternal circulation by IAV infection, IL-6 protein was not detectable from placental lysates and placental and fetal brain *Il6* transcripts were not differentially regulated. Further investigation is needed to discern the impact of heightened maternal IL-6 on fetal development within discrete model systems of MIA, especially when comparing live pathogen infections with bolus administration of LPS, poly I:C, or recombinant IL-6. Overall, it appears the placenta is able to maintain a functional barrier system during moderately severe IAV infection despite indications of mild pathology, proffering protection to the developing fetus.

Only one group, Fatemi and colleagues, has done extensive work in establishing an MIA mouse model of IAV infection by inoculating pregnant mice with human Influenza A/NWS/33 (H1N1) at various gestational time points^4,66–71^. There are several important factors that differ between this model and the one we present here: (a) IAV strain and dose (Fatemi et al. used the H1N1 NWS strain, which possesses the unusual property of being neurotropic in mice, which is not generally a common feature of IAVs; dosing was reported as a dilution of an in-house stock virus and cannot be compared with our study); (b) gestational timing of IAV infection (Fatemi et al. use GD7, 9, 16, and 18); (c) mouse strain (Fatemi et al. use both C57BL/6 and Balb/c); and (d) Fatemi et al.’s use of a broad-spectrum antibiotic (oxytetracycline) in the drinking water to prevent secondary infection^4,66–70^. Overall, this group’s pioneering work revealed the various adverse effects that maternal infection with IAV can have on postnatal mouse offspring brain morphology, chemistry, and transcription, despite the lack of H1N1 viral presence in offspring tissues (^4^ and reviewed in ^17^). In particular, inoculation of pregnant mice at GD9 revealed 38 genes to be dysregulated in the brains of neonatal (P0) progeny^69^, while another group reported 256 differentially expressed genes in the E12.5 embryonic brain using this same IAV model^65^. Here, we found almost no differentially expressed genes in the E17 fetal brain due to maternal IAV X31 infection at GD10. Further investigation into comparable neurodevelopmental and neurocognitive parameters, pre- and postnatally, is needed in order to determine if additional differences exist between the two models. Nevertheless, the lack of transcriptional changes in immune, barrier integrity, and neurodevelopmental genes here suggests that offspring neurological outcomes, if present, are likely to be more mild than what has previously been described in the H1N1 A/NWS/33 infection model^65–68^. Indeed, recent epidemiological data indicates that severe IAV infection during pregnancy significantly elevated the risk of adverse infant outcomes, while less severe infection did not^52^. Our data support a model in which a moderately pathogenic IAV infection is below the severity threshold necessary to induce detrimental effects on fetal brain immune equilibirum.

Given the potential contribution of T_H_17 cell responses in MIA outcomes^72^, one of our major goals was to investigate IL-17 signaling pathways during IAV infection. While respiratory IAV infection increased IL-17A and IL-17C in the lung, IL-17 cytokines were below detectable levels within circulation and did not differ in the intestine. It is possible that this is due to IAV strain differences^73^, as enhanced intestinal T_H_17 cell polarization and IL-17A signaling has been documented in male mice infected with a more pathogenic IAV strain^26^. Even so, this more severe IAV infection did not alter circulating levels of IL-17A^26^, similar to what we report here. This is in stark contrast to poly I:C-induced MIA, wherein increased levels of IL-17A in maternal circulation are readily apparent two days after poly I:C injection^9^. In this model, increased maternal IL-17A augments IL-17RA expression in the fetal brain^9^, whereas our maternal IAV infection model resulted in a minimal but significant downregulation of fetal brain *Il17ra* expression at 7 dpi, which suggests that IL-17A signaling is not pervasive during moderate IAV-induced MIA.

Within the aforementioned poly I:C MIA model, intestinal dendritic cells detect poly I:C directly through TLR3 and induce IL-17A production from pre-existing intestinal T_H_17 cells^20^. Conversely, IAV infection has been shown to indirectly disrupt intestinal microbial communities through lung-derived adaptive responses, which then lead to antimicrobial signaling by intestinal epithelial cells, ultimately promoting T_H_17 cell polarization from resident naive T cells^26^. This is in line with our observation of upregulated *Rorc* within the CD45^+^ intestinal lymphocyte population, suggesting a similar induction of T_H_17 polarization. Importantly, this occurs much later (7 days post IAV inoculation) compared to 12-24 hours post poly I:C injection^9^. Critically, however, it appears that IL-17 cytokine production is restricted to the lung at the timepoint examined in our model, with no detectable changes in IL-17 transcripts or protein in the intestine or in circulation at 7 dpi. Likewise, T_H_17-related gene transcripts (IL-17 receptor *Il17ra* and T_H_17-polarizing *Tgfb1* and *Il23a*) were not differentially regulated within intestinal epithelial cells. While it is possible that changes in IL-17 signaling occurs at time points outside of 7 dpi, these findings bolster the theory that moderate IAV infection may be below severity threshold for eliciting IL-17A responses comparable to levels achieved by poly I:C bolus.

It has previously been shown that a maternal intestinal microbiome that contains segmented filamentous bacteria (SFB) is required for poly I:C-induced offspring abnormalities^20,24^, in line with the particular ability of SFB to promote T_H_17 cell development^25^. Yet IAV infection has been documented to substantially *reduce* SFB numbers, despite concomitant increases in T_H_17 cell polarization and in IL-17A production^26^. Although microbial communities were not investigated here, we did observe a dysregulation in expression of antimicrobial response genes within the intestine, suggesting that endogenous microbes likely play a role in our IAV-induced MIA model. Overall, a marked decrease in colon length of IAV-infected dams (similar to prior observations^26^), in conjunction with differential expression of immune and antimicrobial genes, indicates that even moderately severe IAV infection during pregnancy can disrupt certain components of maternal intestinal health. However, trending increases or stable expression of colonic tight junction proteins, in addition to equivalent circulating levels of LPS-binding protein between IAV-infected and control dams, supports an intact intestinal barrier at this timepoint. Furthermore, the lack of intestinal pathology at 7 dpi across both colonic and ileal H&E-stained sections confirms that overt intestinal injury, outside of colonic shortening, is mostly absent. Overall, these conflicting reports between poly I:C- and IAV-induced intestinal responses epitomize the need for live pathogen MIA models that can further elucidate the influential potential of maternal microbes and adaptive immune responses on offspring outcomes.

Taken together, our data indicate that a clinically relevant and moderately pathogenic IAV infection during pregnancy results in downstream inflammatory sequelea in the placenta but not the fetal brain. While live IAV infection is also characterized by indirect effects within maternal intestinal tissue, IL-17A signaling appears not to play a significant role if the infection is less severe. Critically, our IAV-induced MIA model mirrors a distinct component of epidemiological data wherein only a fraction of infections during pregnancy result in aberrant offspring outcomes, and that offspring psychiatric risk intensifies with increasing infection severity. Overall, our data emphasize the importance of conducting experiments using biologically relevant and translatable MIA models in order to better understand the mechanisms underlying the etiology of neurodevelopmental disorders.

## Supporting information

MIA reporting guidelines checklist

## Acknowledgements

We would like to thank Sydney Schiff, Therese Rajasekera, Zachary Waite, and Hannah Rashidi for their assistance with animal care and tissue processing.

