## Supplementary Figures and Tables for "Moderately pathogenic maternal influenza A virus infection disrupts placental integrity but spares the fetal brain"

Number of Figures: 3

Number of Tables: 7

### Supplementary Figures.

Supplementary Figure S1.

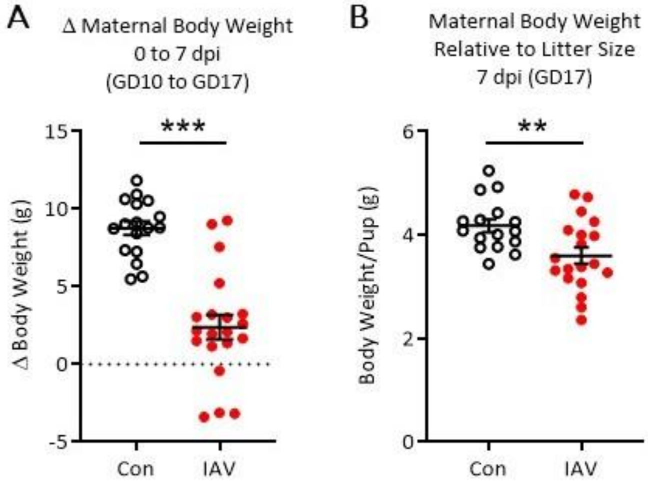


**Supplementary Figure S1. Maternal body mass characteristics. (A)** Pregnant dams administered saline on GD10 (0 dpi) gained at least 5 g body mass by GD17 (7 dpi), while pregnant dams inoculated with IAV gained considerably less weight or lost weight. **(B)** When normalized to litter size ($\frac{\text{body mass (g)}}{\text{number of pups}}$), maternal body weight at GD17 was stunted by IAV-infection; Con: n = 17, IAV: n = 20. IAV = influenza A virus, dpi = days post-inoculation; *** = p < 0.001, ** = p < 0.01.

**Supplementary Figure S2.**

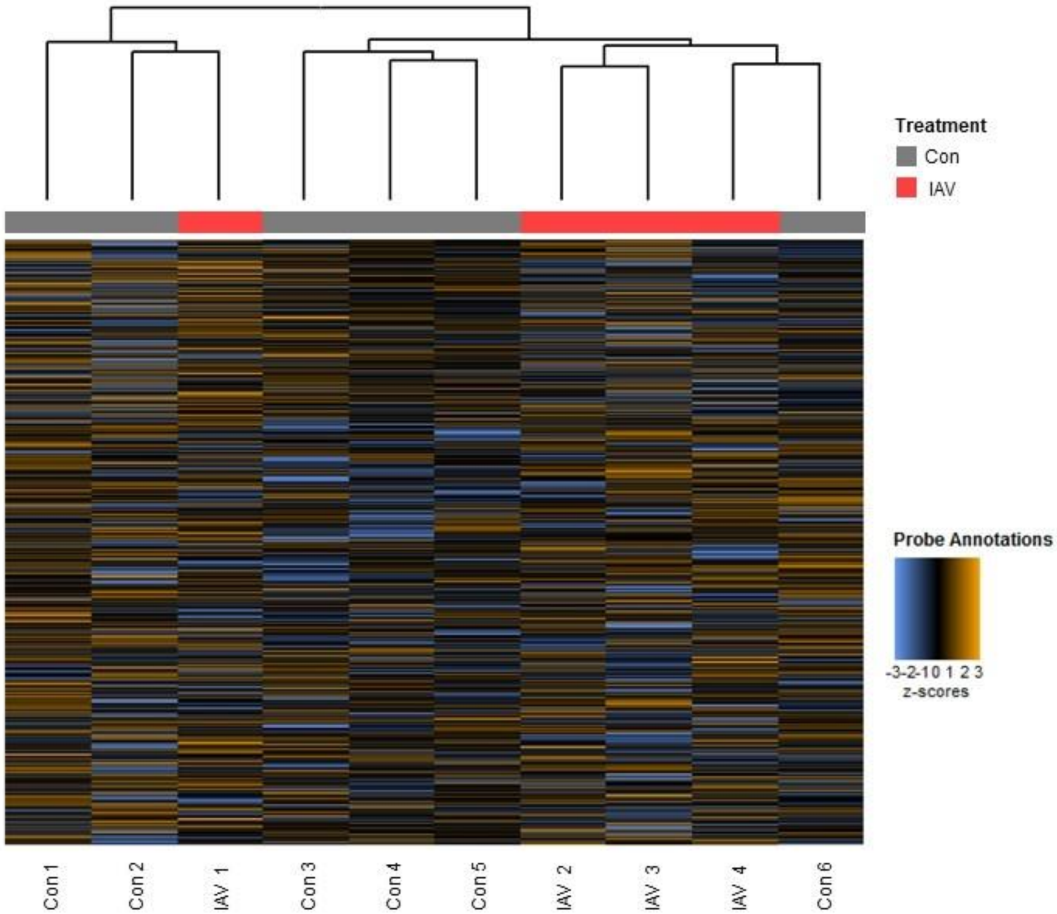


**Supplementary Figure S2. nanoString Neuroinflammation Panel results.** Heatmap of the normalized data, scaled to give all genes equal variance, generated via unsupervised clustering. Orange indicates high expression; blue indicates low expression. Upon FDR correction, there were zero differentially expressed genes due to maternal IAV infection identified within the fetal brain. This plot is meant to provide a high level exploratory view. IAV = influenza A virus.

**Supplementary Figure S3.**

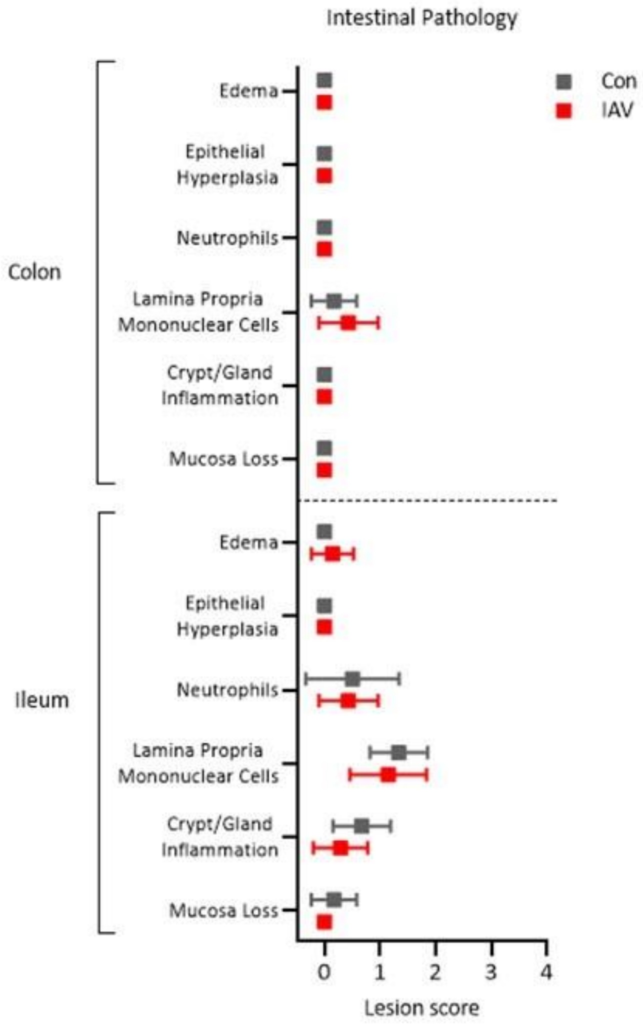


**Supplementary Figure S3. Histopathological scoring of dam intestinal tissue.** Scoring of H&E stained colonic and ileal cross sections revealed no changes across six different categories of inflammatory lesion criteria. The scoring rubric is defined in the Methods. Two sections per dam were scored and averaged; Con: n = 6 dams, IAV: n = 7 dams. IAV = Influenza A virus.

### Supplementary Tables.

Supplemental Table S1. Litter Characteristics.

| Treatment | Control | IAV | p-value |
|---|---|---|---|
| Litter size | 7.5 ± 0.3 | 7.2 ± 0.4 | 0.48 |
| Fetal resorptions | 0.6 ± 0.2 | 0.6 ± 0.3 | 0.77 |

Litter characteristics of gestating dams. Unpaired T tests. Data are presented as mean ± SEM. Control: n = 17, IAV: n = 20.

Supplemental Table S2. Placenta PCR.

| Classification/Function | Gene | Control | IAV | p-value |
|---|---|---|---|---|
| Interleukin-1β; pro-inflammatory | *Il1b* | 1.0 ± 0.10 | 1.01 ± 0.16 | 0.97 |
| Interleukin-6; pro-inflammatory | *Il6* | 1.0 ± 0.05 | 1.22 ± 0.12 | 0.13 |
| Integrin CD11b; complement receptor | *Itgam* | 1.0 ± 0.07 | 0.95 ± 0.15 | 0.76 |
| Tumor necrosis factor; pro-inflammatory | *Tnf* | 1.0 ± 0.06 | 1.04 ± 0.11 | 0.78 |
| Interleukin-17 Receptor A; pro-inflammatory | *Il17ra* | 1.0 ± 0.03 | 1.02 ± 0.05 | 0.71 |
| Type I interferon; anti-viral response | *Ifna4* | 1.0 ± 0.13 | 1.11 ± 0.19 | 0.63 |
| Type I interferon; anti-viral response | *Ifnb1* | 1.0 ± 0.19 | 1.11 ± 0.24 | 0.73 |
| Interferon-induced protein; anti-viral response | *Ifitm3* | 1.0 ± 0.03 | 0.85 ± 0.05 | **0.03** |
| Interferon regulatory factor; anti-viral response | *Irf3* | 1.0 ± 0.06 | 0.95 ± 0.05 | 0.53 |
| Interferon regulatory factor; anti-viral response | *Irf7* | 1.0 ± 0.04 | 0.92 ± 0.10 | 0.44 |
| Recognition and uptake of bacterial components | *Tlr2* | 1.0 ± 0.08 | 0.89 ± 0.13 | 0.45 |
| Recognition and uptake of bacterial components | *Tlr4* | 1.0 ± 0.05 | 0.97 ± 0.15 | 0.81 |
| Antimicrobial adaptor protein; pro-inflammatory | *Myd88* | 1.0 ± 0.06 | 1.00 ± 0.08 | 0.97 |
| Recognition and uptake of bacterial components | *Ptafr* | 1.0 ± 0.10 | 1.14 ± 0.09 | 0.37 |
| Interleukin-22; antimicrobial defense | *Il22* | 1.0 ± 0.17 | 1.23 ± 0.18 | 0.38 |
| Bactericidal C-type lectin; antimicrobial | *Reg3g* | 1.0 ± 0.30 | 1.49 ± 0.73 | 0.60 |
| Alpha Defensin 1; antimicrobial | *Defa1* | 1.0 ± 0.14 | 1.12 ± 0.08 | 0.44 |
| Tight junction protein; barrier integrity | *Cldn1* | 1.0 ± 0.05 | 1.17 ± 0.07 | *0.08* |
| Tight junction protein; barrier integrity | *Cldn2* | 1.0 ± 0.12 | 1.43 ± 0.28 | 0.20 |
| Tight junction protein; barrier integrity | *Cldn5* | 1.0 ± 0.08 | 0.92 ± 0.05 | 0.46 |
| Tight junction protein; barrier integrity | *Ocln* | 1.0 ± 0.08 | 1.14 ± 0.17 | 0.50 |
| Matrix metalloproteinase 2; ECM remodeling | *Mmp2* | 1.0 ± 0.05 | 0.85 ± 0.06 | *0.07* |
| Matrix metalloproteinase 9; ECM remodeling | *Mmp9* | 1.0 ± 0.06 | 1.20 ± 0.14 | 0.23 |

Relative expression of 23 different immune and barrier integrity genes in placental tissue of litters from saline control and IAV-infected gestating dams at GD17 (1 placenta/litter). Data are presented as mean ± SEM; n = 6-13/group. Bold font = $p < 0.05$, italicized font = $p < 0.10$. ECM = extracellular matrix.

Supplemental Table S3. Uterine PCR.

| Classification/Function | Gene | Control | IAV | p-value |
|---|---|---|---|---|
| Granzyme B; immune-cell driven apoptosis | *Gzmb* | 1.0 ± 0.18 | 1.01 ± 0.33 | 0.98 |
| Type I interferon; anti-viral response | *Ifna4* | 1.0 ± 0.24 | 2.03 ± 0.74 | 0.16 |
| Type I interferon; anti-viral response | *Ifnb1* | 1.0 ± 0.22 | 1.41 ± 0.43 | 0.38 |
| Type II interferon; anti-viral response | *Ifng* | 1.0 ± 0.08 | 1.03 ± 0.12 | 0.85 |
| Bactericidal C-type lectin; antimicrobial | *Reg3b* | 1.0 ± 0.14 | 1.77 ± 0.58 | 0.19 |
| Bactericidal C-type lectin; antimicrobial | *Reg3g* | 1.0 ± 0.35 | 0.86 ± 0.23 | 0.76 |
| Interleukin-10; anti-inflammatory | *Il10* | 1.0 ± 0.05 | 1.08 ± 0.23 | 0.72 |
| Tumor necrosis factor; pro-inflammatory | *Tnf* | 1.0 ± 0.08 | 1.01 ± 0.12 | 0.97 |

Relative expression of immune genes in uterine tissue of saline control and IAV-infected gestating dams at GD17. Data are presented as mean ± SEM; n = 6-8/group.

**Supplemental Table S4. Fetal brain differential gene expression data from the nanoString nCounter Mouse Neuroinflammation Panel –** Presented as a separate file.

Supplemental Table S5. Fetal brain PCR.

| Classification/Function | Gene | Control | IAV | p-value |
|---|---|---|---|---|
| Tumor necrosis factor; pro-inflammatory | *Tnf* | 1.0 ± 0.11 | 0.91 ± 0.08 | 0.55 |
| Interleukin-6; pro-inflammatory | *Il6* | 1.0 ± 0.08 | 1.03 ± 0.16 | 0.86 |
| Integrin CD11b; complement receptor | *Itgam* | 1.0 ± 0.06 | 0.96 ± 0.04 | 0.67 |
| Interleukin-17 Receptor A; pro-inflammatory | *Il17ra* | 1.0 ± 0.04 | 0.85 ± 0.02 | **0.009** |
| Interleukin-1 beta; pro-inflammatory | *Il1b* | 1.0 ± 0.16 | 0.57 ± 0.04 | **0.046** |
| Type I interferon; anti-viral response | *Ifna4* | 1.0 ± 0.23 | 0.94 ± 0.14 | 0.85 |
| Type I interferon; anti-viral response | *Ifnb1* | 1.0 ± 0.22 | 0.87 ± 0.15 | 0.66 |
| Type II interferon; anti-viral response | *Ifng* | 1.0 ± 0.11 | 1.45 ± 0.42 | 0.26 |
| Interferon-induced protein; anti-viral response | *Ifit1* | 1.0 ± 0.24 | 0.46 ± 0.10 | 0.12 |
| Interferon-induced protein; anti-viral response | *Ifitm3* | 1.0 ± 0.10 | 0.99 ± 0.08 | 0.95 |
| Interferon regulatory factor; anti-viral response | *Irf3* | 1.0 ± 0.04 | 0.90 ± 0.03 | *0.08* |
| Interferon regulatory factor; anti-viral response | *Irf7* | 1.0 ± 0.17 | 0.99 ± 0.26 | 0.97 |
| Recognition and uptake of bacterial components | *Tlr2* | 1.0 ± 0.12 | 0.86 ± 0.12 | 0.42 |
| Recognition and uptake of bacterial components | *Tlr4* | 1.0 ± 0.13 | 0.85 ± 0.10 | 0.42 |
| Recognition and uptake of bacterial components | *Ptafr* | 1.0 ± 0.11 | 0.94 ± 0.08 | 0.70 |
| Tight junction protein; barrier integrity | *Cldn1* | 1.0 ± 0.18 | 0.92 ± 0.13 | 0.74 |
| Tight junction protein; barrier integrity | *Cldn2* | 1.0 ± 0.23 | 0.86 ± 0.12 | 0.64 |
| Tight junction protein; barrier integrity | *Cldn5* | 1.0 ± 0.11 | 0.82 ± 0.09 | 0.26 |
| Tight junction protein; barrier integrity | *Ocln* | 1.0 ± 0.18 | 0.82 ± 0.11 | 0.43 |
| Transforming growth factor; neuroprotective | *Tgfb1* | 1.0 ± 0.10 | 0.81 ± 0.09 | 0.21 |
| Neurotrophic factor; neuroprotective | *Bdnf* | 1.0 ± 0.06 | 0.99 ± 0.08 | 0.90 |
| Purinergic receptor; microglia responses | *P2ry12* | 1.0 ± 0.05 | 1.04 ± 0.06 | 0.63 |
| Fractalkine; microglia-neuron interactions | *Cx3cl1* | 1.0 ± 0.08 | 0.93 ± 0.11 | 0.60 |

Relative expression of 23 different immune, barrier integrity, and neurodevelopmental genes in fetal brain tissue of litters from saline control and IAV-infected gestating dams at GD17 (1 fetal brain/litter). Data are presented as mean ± SEM; n = 6-8/group. Bold font = $p < 0.05$, italicized font = $p < 0.10$.

Supplemental Table S6. Small intestine intraepithelial lymphocyte (IEL) PCR.

| Classification/Function | Gene | Control | IAV | p-value |
|---|---|---|---|---|
| Type I interferon; anti-viral response | *Ifna4* | 1.0 ± 0.32 | 0.50 ± 0.13 | 0.32 |
| Type I interferon; anti-viral response | *Ifnb1* | 1.0 ± 0.27 | 0.44 ± 0.15 | 0.20 |
| Type II interferon; anti-viral response | *Ifng* | 1.0 ± 0.21 | 1.00 ± 0.18 | 0.99 |
| Interleukin-17; pro-inflammatory | *Il17* | 1.0 ± 0.23 | 1.54 ± 0.71 | 0.43 |
| Interleukin-17 Receptor A; pro-inflammatory | *Il17ra* | 1.0 ± 0.05 | 0.99 ± 0.03 | 0.89 |
| Interleukin-6; pro-inflammatory | *Il6* | 1.0 ± 0.27 | 1.45 ± 0.30 | 0.30 |

Relative expression of immune genes in the $CD45^+$ population (IEL) of small intestinal tissue from saline control and IAV-infected gestating dams at GD17. Data are presented as mean ± SEM; n = 6-8/group.

Supplementary Table S7. Gene Primers.

| Gene | Assay ID[a] |
|---|---|
| *Bdnf* | Mm00432069_m1 |
| *Ccl2* | Mm00441242_m1 |
| *Cldn1* | Mm00516701_m1 |
| *Cldn2* | Mm00516703_s1 |
| *Cldn5* | Mm00727012_s1 |
| *Cx3cl1* | Mm00436454_m1 |
| *Defa1* | Mm02524428_g1 |
| *Gzmb* | Mm00442837_m1 |
| *Ifit1* | Mm00515153_m1 |
| *Ifitm3* | Mm00847057_s1 |
| *Ifna4* | Mm00833969_s1 |
| *Ifnb1* | Mm00439552_s1 |
| *Ifng* | Mm01168134_m1 |
| *Il10* | Mm01288386_m1 |
| *Il17a* | Mm00439618_m1 |
| *Il17ra* | Mm00434214_m1 |
| *Il1b* | Mm00434228_m1 |
| *Il22* | Mm01226722_g1 |
| *Il6* | Mm00446190_m1 |
| *Irf3* | Mm00516784_m1 |
| *Irf7* | Mm00516793_g1 |
| *Itgam* | Mm00434455_m1 |
| *Lbp* | Mm00493139_m1 |
| *Mmp2* | Mm00439498_m1 |
| *Mmp9* | Mm00442991_m1 |
| *Myd88* | Mm00440338_m1 |
| *Ocln* | Mm00500912_m1 |
| *P2ry12* | Mm00446026_m1 |
| *Ptafr* | Mm01187337_m1 |
| *Reg3b* | Mm00440616_g1 |
| *Reg3g* | Mm00441127_m1 |
| *Rorc* | Mm01261022_m1 |
| *Rpl19* | Mm02601633_g1 |
| *Tgfb1* | Mm01178820_m1 |
| *Tlr2* | Mm01213946_g1 |
| *Tlr4* | Mm00445273_m1 |
| *Tnf* | Mm00443258_m1 |

[a] Applied Biosystems TaqMan Gene Expression Assay identification number.
