## Supplemental Table S4 for "Moderately pathogenic maternal influenza A virus infection disrupts placental integrity but spares the fetal brain"

Supplemental Table S4. Fetal brain differential gene expression data from the nanoString nCounter Mouse Neuroinflammation Panel.

| mRNA | Log2 fold change | Std error (log2) | Lower confidence limit (log2) | Upper confidence limit (log2) | Unadj p-value | Bonferroni adj p-value | Method | Probe ID |
|---|---|---|---|---|---|---|---|---|
| **Jun** | -0.202 | 0.0328 | -0.266 | -0.137 | 0.000275 | 0.162 | loglinear | NM_010591.2:2212 |
| **Rpl28** | -0.067 | 0.0137 | -0.0939 | -0.0402 | 0.0012 | 0.712 | loglinear | NM_009081.2:106 |
| **S1pr3** | -0.324 | 0.0677 | -0.457 | -0.191 | 0.00137 | 0.811 | loglinear | NM_010101.3:2939 |
| **Hmgb1** | -0.169 | 0.036 | -0.239 | -0.0982 | 0.00157 | 0.927 | loglinear | NM_010439.3:1574 |
| **Ung** | -0.383 | 0.0852 | -0.551 | -0.216 | 0.00201 | 1 | loglinear | NM_001040691.1:336 |
| **Msn** | -0.439 | 0.102 | -0.638 | -0.239 | 0.00257 | 1 | loglinear | NM_010833.2:515 |
| **Rps3** | 0.172 | 0.0409 | 0.0918 | 0.252 | 0.00297 | 1 | loglinear | NM_012052.2:886 |
| **Tbr1** | -0.36 | 0.086 | -0.529 | -0.192 | 0.00305 | 1 | loglinear | NM_009322.3:2354 |
| **Lag3** | 0.772 | 0.179 | 0.42 | 1.12 | 0.00357 | 1 | Wald | NM_008479.1:1700 |
| **Hps4** | -0.344 | 0.0887 | -0.518 | -0.17 | 0.00466 | 1 | loglinear | NM_138646.3:1265 |
| **Casp3** | 0.116 | 0.0304 | 0.0567 | 0.176 | 0.00505 | 1 | loglinear | NM_009810.2:630 |
| **Crip1** | -0.287 | 0.0858 | -0.455 | -0.119 | 0.0101 | 1 | loglinear | NM_007763.3:47 |
| **Braf** | 0.121 | 0.0372 | 0.0485 | 0.194 | 0.0115 | 1 | loglinear | NM_139294.5:1102 |
| **Zfp367** | -0.264 | 0.0811 | -0.423 | -0.105 | 0.0117 | 1 | loglinear | NM_175494.4:1235 |
| **Tnfrsf1a** | -0.364 | 0.116 | -0.592 | -0.137 | 0.0138 | 1 | loglinear | NM_011609.2:615 |
| **Ptms** | 0.0859 | 0.0275 | 0.0319 | 0.14 | 0.0142 | 1 | loglinear | NM_026988.2:755 |
| **Cx3cr1** | -0.264 | 0.0848 | -0.431 | -0.0983 | 0.0142 | 1 | loglinear | NM_009987.3:2696 |
| **Prkar2a** | 0.0962 | 0.0324 | 0.0327 | 0.16 | 0.0179 | 1 | loglinear | NM_008924.2:2135 |
| **Pak1** | 0.0756 | 0.0257 | 0.0253 | 0.126 | 0.0186 | 1 | loglinear | NM_011035.2:1615 |
| **Mapk14** | 0.11 | 0.0375 | 0.0363 | 0.183 | 0.019 | 1 | loglinear | NM_011951.2:1420 |
| **Hells** | -0.223 | 0.0765 | -0.373 | -0.0727 | 0.0196 | 1 | loglinear | NM_008234.3:1082 |
| **Suz12** | 0.0537 | 0.0187 | 0.0172 | 0.0903 | 0.0205 | 1 | loglinear | NM_199196.1:820 |
| **Ms4a4a** | -0.648 | 0.219 | -1.08 | -0.219 | 0.021 | 1 | Wald | XM_003086124.1:252 |
| **Npnt** | 0.2 | 0.0703 | 0.0625 | 0.338 | 0.0215 | 1 | loglinear | NM_001029836.1:2650 |
| **Ifnar1** | 0.116 | 0.041 | 0.036 | 0.197 | 0.0219 | 1 | loglinear | NM_010508.1:1195 |
| **Xiap** | 0.0974 | 0.0346 | 0.0295 | 0.165 | 0.0228 | 1 | loglinear | NM_009688.2:1654 |
| **Tie1** | -0.236 | 0.0851 | -0.403 | -0.0696 | 0.024 | 1 | loglinear | NM_011587.2:2715 |
| **Serping1** | -0.249 | 0.0903 | -0.426 | -0.0724 | 0.0246 | 1 | loglinear | NM_009776.3:1480 |

| | | | | | | | | |
|---|---|---|---|---|---|---|---|---|
| **Pink1** | 0.186 | 0.0674 | 0.0536 | 0.318 | 0.0249 | 1 | loglinear | NM_026880.2:688 |
| **Dna2** | -0.513 | 0.186 | -0.878 | -0.147 | 0.025 | 1 | lm.nb | NM_177372.3:1734 |
| **Atg3** | 0.101 | 0.0376 | 0.0273 | 0.175 | 0.0277 | 1 | loglinear | NM_026402.3:862 |
| **Rps21** | -0.0728 | 0.0273 | -0.126 | -0.0194 | 0.0283 | 1 | loglinear | NM_025587.2:153 |
| **Rac1** | 0.0501 | 0.0196 | 0.0116 | 0.0886 | 0.0342 | 1 | loglinear | NM_009007.2:1045 |
| **Stat1** | -0.384 | 0.149 | -0.676 | -0.0919 | 0.0367 | 1 | Wald | NM_009283.3:1590 |
| **Traf2** | -0.21 | 0.084 | -0.374 | -0.0451 | 0.0371 | 1 | loglinear | NM_009422.2:1334 |
| **Vps4b** | 0.167 | 0.0678 | 0.0336 | 0.3 | 0.0396 | 1 | loglinear | NM_009190.2:640 |
| **Xrcc6** | 0.219 | 0.0912 | 0.0399 | 0.398 | 0.0433 | 1 | loglinear | NM_010247.2:1640 |
| **H2afx** | -0.17 | 0.0717 | -0.311 | -0.0297 | 0.045 | 1 | loglinear | NM_010436.2:980 |
| **Itgb5** | -0.139 | 0.0595 | -0.255 | -0.0222 | 0.0479 | 1 | loglinear | NM_001145884.1:1270 |
| **Srgn** | -0.342 | 0.143 | -0.622 | -0.0606 | 0.0487 | 1 | Wald | NM_011157.2:168 |
| **Atm** | -0.135 | 0.058 | -0.248 | -0.021 | 0.0488 | 1 | loglinear | NM_007499.2:5543 |
| **Axl** | -0.207 | 0.0914 | -0.386 | -0.0278 | 0.0533 | 1 | loglinear | NM_009465.3:3820 |
| **Ctss** | 0.305 | 0.132 | 0.0474 | 0.563 | 0.0534 | 1 | Wald | NM_021281.2:740 |
| **Itga7** | -0.378 | 0.163 | -0.698 | -0.0579 | 0.0538 | 1 | Wald | NM_008398.2:2435 |
| **Cd47** | 0.0997 | 0.0445 | 0.0125 | 0.187 | 0.0553 | 1 | loglinear | NM_010581.3:165 |
| **Il10rb** | -0.298 | 0.136 | -0.564 | -0.0319 | 0.0595 | 1 | loglinear | NM_008349.5:465 |
| **Rad51** | -0.307 | 0.143 | -0.587 | -0.0265 | 0.0642 | 1 | loglinear | NM_011234.4:286 |
| **Hif1a** | 0.0545 | 0.0262 | 0.00319 | 0.106 | 0.0709 | 1 | loglinear | NM_010431.2:1294 |
| **Mapk10** | 0.0987 | 0.0487 | 0.00328 | 0.194 | 0.0772 | 1 | loglinear | NM_001081567.1:1496 |
| **Slc17a6** | 0.147 | 0.0737 | 0.0027 | 0.292 | 0.0809 | 1 | loglinear | NM_080853.3:2825 |
| **C1qa** | -0.299 | 0.15 | -0.594 | -0.00507 | 0.0813 | 1 | loglinear | NM_007572.2:566 |
| **Islr2** | -0.0736 | 0.0372 | -0.147 | -0.000706 | 0.0832 | 1 | loglinear | NM_001161538.1:1782 |
| **Kdm2a** | 0.0626 | 0.0317 | 0.000541 | 0.125 | 0.0834 | 1 | loglinear | NM_001001984.2:4160 |
| **Plekhb1** | 0.255 | 0.126 | 0.00723 | 0.503 | 0.0835 | 1 | Wald | NM_001163184.1:1616 |
| **Hat1** | -0.148 | 0.0762 | -0.298 | 0.000992 | 0.0874 | 1 | loglinear | NM_026115.4:1270 |
| **Ccni** | -0.0589 | 0.0303 | -0.118 | 0.000505 | 0.0879 | 1 | loglinear | NM_017367.3:1180 |
| **Dock2** | -0.434 | 0.219 | -0.864 | -0.00353 | 0.0887 | 1 | Wald | NM_033374.3:2410 |
| **Pik3ca** | 0.0755 | 0.0391 | -0.00119 | 0.152 | 0.0898 | 1 | loglinear | NM_008839.1:1255 |
| **Akt2** | -0.186 | 0.0965 | -0.375 | 0.00326 | 0.0902 | 1 | loglinear | NM_001110208.1:2504 |
| **Tfg** | 0.0499 | 0.0261 | -0.0013 | 0.101 | 0.0925 | 1 | loglinear | NM_001252443.1:578 |

| | | | | | | | | |
|---|---|---|---|---|---|---|---|---|
| **Plp1** | -0.3 | 0.158 | -0.61 | 0.0107 | 0.095 | 1 | lm.nb | NM_011123.2:795 |
| **Fcgr2b** | -0.38 | 0.199 | -0.77 | 0.00978 | 0.0976 | 1 | Wald | NM_001077189.1:1225 |
| **Trp53bp2** | -0.127 | 0.0683 | -0.261 | 0.0069 | 0.1 | 1 | loglinear | NM_173378.2:2328 |
| **Ncor2** | -0.0887 | 0.0484 | -0.183 | 0.00607 | 0.104 | 1 | loglinear | NM_011424.2:1156 |
| **Lgmn** | 0.158 | 0.0873 | -0.0127 | 0.329 | 0.107 | 1 | loglinear | NM_011175.3:370 |
| **Irf7** | -0.478 | 0.261 | -0.99 | 0.0339 | 0.11 | 1 | Wald | NM_016850.2:705 |
| **Nrm** | -0.129 | 0.072 | -0.271 | 0.0117 | 0.11 | 1 | loglinear | NM_134122.2:466 |
| **Nbn** | 0.167 | 0.0938 | -0.0169 | 0.351 | 0.113 | 1 | loglinear | NM_013752.3:1000 |
| **Ifnar2** | -0.138 | 0.0775 | -0.289 | 0.0144 | 0.114 | 1 | loglinear | NM_001110498.1:725 |
| **Sesn1** | -0.112 | 0.0636 | -0.237 | 0.0123 | 0.115 | 1 | loglinear | NM_001013370.2:497 |
| **Fabp5** | 0.0528 | 0.03 | -0.00604 | 0.112 | 0.117 | 1 | loglinear | NM_010634.3:430 |
| **Creb1** | -0.0558 | 0.0318 | -0.118 | 0.00658 | 0.118 | 1 | loglinear | NM_001037726.1:2734 |
| **Sall1** | -0.217 | 0.126 | -0.464 | 0.0291 | 0.122 | 1 | lm.nb | NM_021390.3:4875 |
| **Hdac1** | -0.164 | 0.0954 | -0.351 | 0.0226 | 0.123 | 1 | loglinear | NM_008228.2:470 |
| **Dapk1** | -0.083 | 0.0482 | -0.177 | 0.0115 | 0.123 | 1 | loglinear | NM_134062.1:4935 |
| **Rps9** | -0.056 | 0.0326 | -0.12 | 0.00796 | 0.125 | 1 | loglinear | NM_029767.2:173 |
| **Apex1** | -0.0765 | 0.0449 | -0.164 | 0.0115 | 0.127 | 1 | loglinear | NM_009687.2:289 |
| **Aldh1l1** | 0.185 | 0.109 | -0.0282 | 0.398 | 0.127 | 1 | loglinear | NM_027406.1:1340 |
| **Ppp3cb** | 0.0725 | 0.0427 | -0.0112 | 0.156 | 0.128 | 1 | loglinear | NM_008914.2:2950 |
| **Bbc3** | 0.211 | 0.125 | -0.0336 | 0.457 | 0.129 | 1 | loglinear | NM_133234.1:1461 |
| **Fancd2** | -0.283 | 0.165 | -0.606 | 0.0408 | 0.13 | 1 | Wald | NM_001033244.3:1146 |
| **Traf3** | 0.113 | 0.0676 | -0.0191 | 0.246 | 0.132 | 1 | loglinear | NM_011632.3:884 |
| **Cdk20** | -0.234 | 0.14 | -0.507 | 0.0401 | 0.133 | 1 | lm.nb | NM_053180.2:352 |
| **Mcm2** | -0.233 | 0.143 | -0.513 | 0.0465 | 0.141 | 1 | lm.nb | NM_008564.2:2585 |
| **Abcc8** | -0.253 | 0.153 | -0.553 | 0.0471 | 0.142 | 1 | Wald | NM_011510.3:1740 |
| **Hdac4** | 0.0815 | 0.0503 | -0.0171 | 0.18 | 0.144 | 1 | loglinear | NM_207225.1:2800 |
| **Lsr** | -0.135 | 0.0837 | -0.299 | 0.0288 | 0.145 | 1 | loglinear | NM_001164184.1:445 |
| **Rps10** | -0.0719 | 0.0446 | -0.159 | 0.0156 | 0.146 | 1 | loglinear | NM_025963.3:318 |
| **Plxnb3** | 0.339 | 0.207 | -0.0673 | 0.745 | 0.146 | 1 | Wald | NM_019587.2:2862 |
| **Cdc25a** | -0.131 | 0.0812 | -0.29 | 0.0285 | 0.146 | 1 | loglinear | NM_007658.3:855 |
| **Hpgds** | 0.255 | 0.157 | -0.0524 | 0.563 | 0.148 | 1 | Wald | NM_019455.4:425 |
| **Crem** | 0.202 | 0.126 | -0.0454 | 0.449 | 0.148 | 1 | loglinear | NM_001110853.1:1840 |

| | | | | | | | | |
|---|---|---|---|---|---|---|---|---|
| **Ikbkg** | -0.159 | 0.0996 | -0.354 | 0.0365 | 0.15 | 1 | loglinear | NM_178590.2:525 |
| **Brd2** | 0.0601 | 0.0378 | -0.014 | 0.134 | 0.151 | 1 | loglinear | NM_010238.3:2800 |
| **Top2a** | -0.264 | 0.168 | -0.593 | 0.0652 | 0.155 | 1 | lm.nb | NM_011623.2:1953 |
| **Gria4** | 0.134 | 0.0853 | -0.0334 | 0.301 | 0.156 | 1 | loglinear | NM_001113180.1:1274 |
| **Akt1** | -0.0658 | 0.0422 | -0.148 | 0.0168 | 0.157 | 1 | loglinear | NM_001165894.1:898 |
| **Egfr** | 0.351 | 0.225 | -0.0909 | 0.792 | 0.158 | 1 | lm.nb | NM_207655.2:1335 |
| **Bax** | 0.0876 | 0.0563 | -0.0228 | 0.198 | 0.158 | 1 | loglinear | NM_007527.3:735 |
| **Nfkb2** | 0.292 | 0.186 | -0.0731 | 0.658 | 0.161 | 1 | Wald | NM_019408.2:1150 |
| **Parp2** | 0.108 | 0.0702 | -0.0298 | 0.245 | 0.163 | 1 | loglinear | NM_009632.2:1325 |
| **Kcnj10** | 0.156 | 0.102 | -0.0446 | 0.356 | 0.166 | 1 | loglinear | NM_001039484.1:400 |
| **Mdc1** | -0.0812 | 0.0535 | -0.186 | 0.0236 | 0.167 | 1 | loglinear | NM_001010833.2:5900 |
| **Rela** | -0.136 | 0.0899 | -0.313 | 0.0398 | 0.168 | 1 | loglinear | NM_009045.4:645 |
| **Il1rl2** | 0.362 | 0.236 | -0.0999 | 0.824 | 0.168 | 1 | Wald | NM_133193.3:860 |
| **Myc** | 0.0777 | 0.0515 | -0.0232 | 0.179 | 0.17 | 1 | loglinear | NM_010849.4:630 |
| **Lair1** | 0.272 | 0.179 | -0.0784 | 0.623 | 0.172 | 1 | Wald | NM_001113474.1:1865 |
| **Arc** | 0.323 | 0.214 | -0.0961 | 0.741 | 0.175 | 1 | Wald | NM_018790.2:2715 |
| **Kit** | 0.182 | 0.124 | -0.06 | 0.425 | 0.179 | 1 | loglinear | NM_001122733.1:4275 |
| **Fgf13** | 0.101 | 0.0687 | -0.0334 | 0.236 | 0.179 | 1 | loglinear | NM_010200.2:700 |
| **Hdac2** | -0.0619 | 0.042 | -0.144 | 0.0205 | 0.179 | 1 | loglinear | NM_008229.2:1010 |
| **Chn2** | -0.109 | 0.0751 | -0.256 | 0.0379 | 0.184 | 1 | loglinear | NM_001163640.1:1510 |
| **Setd1a** | -0.132 | 0.0909 | -0.31 | 0.0461 | 0.184 | 1 | loglinear | NM_178029.3:1612 |
| **Hus1** | -0.172 | 0.12 | -0.407 | 0.0627 | 0.189 | 1 | loglinear | NM_008316.2:2505 |
| **Mertk** | 0.215 | 0.148 | -0.0754 | 0.506 | 0.19 | 1 | Wald | NM_008587.1:1320 |
| **Cd83** | 0.21 | 0.147 | -0.0783 | 0.498 | 0.191 | 1 | lm.nb | NM_009856.2:1624 |
| **Pex14** | 0.0972 | 0.0684 | -0.0368 | 0.231 | 0.193 | 1 | loglinear | NM_019781.2:597 |
| **Dst** | -0.101 | 0.0712 | -0.241 | 0.0385 | 0.194 | 1 | loglinear | NM_010081.2:226 |
| **Csk** | -0.0783 | 0.0555 | -0.187 | 0.0303 | 0.195 | 1 | loglinear | NM_007783.2:268 |
| **Rad51b** | -0.34 | 0.239 | -0.808 | 0.128 | 0.198 | 1 | Wald | NM_009014.3:340 |
| **Birc2** | 0.0593 | 0.0424 | -0.0239 | 0.143 | 0.2 | 1 | loglinear | NM_007465.2:1230 |
| **Itga6** | 0.061 | 0.0437 | -0.0247 | 0.147 | 0.201 | 1 | loglinear | NM_008397.3:910 |
| **Plxdc2** | 0.111 | 0.0804 | -0.0465 | 0.269 | 0.204 | 1 | loglinear | NM_026162.5:1580 |
| **Rad51c** | -0.203 | 0.146 | -0.489 | 0.0837 | 0.208 | 1 | Wald | NM_053269.3:402 |

| | | | | | | | | |
|---|---|---|---|---|---|---|---|---|
| **Prkar1a** | 0.0541 | 0.0396 | -0.0235 | 0.132 | 0.209 | 1 | loglinear | NM_021880.2:2235 |
| **Ehmt2** | 0.0378 | 0.0277 | -0.0166 | 0.0922 | 0.21 | 1 | loglinear | NM_145830.1:3475 |
| **Topbp1** | -0.0807 | 0.0594 | -0.197 | 0.0357 | 0.211 | 1 | loglinear | NM_176979.5:3564 |
| **Casp7** | 0.0936 | 0.0697 | -0.0429 | 0.23 | 0.216 | 1 | loglinear | NM_007611.2:1468 |
| **Bnip3l** | 0.0653 | 0.0487 | -0.0302 | 0.161 | 0.217 | 1 | loglinear | NM_009761.3:1738 |
| **Rad17** | -0.0995 | 0.0749 | -0.246 | 0.0473 | 0.221 | 1 | loglinear | NM_001044371.1:386 |
| **Pecam1** | -0.148 | 0.112 | -0.368 | 0.0717 | 0.223 | 1 | loglinear | NM_008816.2:1100 |
| **Rad50** | -0.09 | 0.0681 | -0.224 | 0.0436 | 0.223 | 1 | loglinear | NM_009012.2:4165 |
| **Vps4a** | 0.0967 | 0.0735 | -0.0472 | 0.241 | 0.224 | 1 | loglinear | NM_126165.1:994 |
| **Coa5** | -0.0572 | 0.0434 | -0.142 | 0.0279 | 0.224 | 1 | loglinear | NM_198006.4:1808 |
| **Myrf** | 0.218 | 0.165 | -0.105 | 0.541 | 0.228 | 1 | Wald | NM_001033481.1:4465 |
| **Smc1a** | -0.0618 | 0.0476 | -0.155 | 0.0314 | 0.23 | 1 | loglinear | NM_019710.2:1675 |
| **Eomes** | -0.27 | 0.209 | -0.681 | 0.14 | 0.233 | 1 | lm.nb | NM_010136.2:2665 |
| **Atp6v0e** | -0.0883 | 0.0685 | -0.223 | 0.0459 | 0.233 | 1 | loglinear | NM_025272.2:585 |
| **Dlg1** | -0.0825 | 0.064 | -0.208 | 0.0429 | 0.233 | 1 | loglinear | NM_001252433.1:1064 |
| **Slc2a1** | -0.11 | 0.0866 | -0.28 | 0.0593 | 0.238 | 1 | loglinear | NM_011400.3:2190 |
| **Casp2** | -0.0978 | 0.0775 | -0.25 | 0.0542 | 0.243 | 1 | loglinear | NM_007610.1:420 |
| **Bmi1** | -0.101 | 0.0798 | -0.257 | 0.0558 | 0.243 | 1 | loglinear | NM_007552.4:3354 |
| **Vegfa** | 0.11 | 0.0871 | -0.0611 | 0.28 | 0.244 | 1 | loglinear | NM_001025250.3:3015 |
| **P2rx7** | -0.166 | 0.132 | -0.424 | 0.0935 | 0.251 | 1 | Wald | NM_001038839.2:378 |
| **Lamp1** | 0.0624 | 0.0507 | -0.037 | 0.162 | 0.253 | 1 | loglinear | NM_010684.2:2080 |
| **Ttr** | -0.267 | 0.217 | -0.692 | 0.158 | 0.253 | 1 | lm.nb | NM_013697.4:855 |
| **AI464131** | 0.203 | 0.166 | -0.122 | 0.527 | 0.256 | 1 | lm.nb | NM_001085515.2:1232 |
| **Dlx1** | -0.155 | 0.127 | -0.403 | 0.0933 | 0.256 | 1 | lm.nb | NM_010053.1:2052 |
| **Cdkn1a** | 0.159 | 0.131 | -0.0981 | 0.416 | 0.26 | 1 | loglinear | NM_007669.4:1670 |
| **Kmt2c** | -0.0521 | 0.0431 | -0.137 | 0.0324 | 0.261 | 1 | loglinear | NM_001081383.1:7075 |
| **Rgl1** | -0.154 | 0.126 | -0.401 | 0.0932 | 0.262 | 1 | Wald | NM_016846.3:3320 |
| **Tgfb1** | 0.172 | 0.141 | -0.105 | 0.449 | 0.262 | 1 | Wald | NM_011577.1:1470 |
| **Serpinf1** | -0.271 | 0.225 | -0.713 | 0.171 | 0.263 | 1 | lm.nb | NM_011340.3:745 |
| **Dlx2** | -0.166 | 0.139 | -0.439 | 0.106 | 0.266 | 1 | loglinear | NM_010054.2:1891 |
| **Slc1a3** | 0.11 | 0.0923 | -0.071 | 0.291 | 0.268 | 1 | loglinear | NM_148938.3:3865 |
| **Ppfia4** | 0.158 | 0.133 | -0.102 | 0.418 | 0.268 | 1 | lm.nb | NM_001144855.1:454 |

| | | | | | | | | |
|---|---|---|---|---|---|---|---|---|
| **Lacc1** | 0.201 | 0.168 | -0.127 | 0.53 | 0.268 | 1 | Wald | NM_172488.2:1025 |
| **Dock1** | -0.114 | 0.0962 | -0.302 | 0.0749 | 0.271 | 1 | loglinear | NM_001033420.2:405 |
| **Arid1a** | -0.0599 | 0.0508 | -0.159 | 0.0396 | 0.272 | 1 | loglinear | NM_001080819.1:5193 |
| **Psmb8** | 0.262 | 0.22 | -0.169 | 0.694 | 0.272 | 1 | Wald | NM_010724.2:362 |
| **Tmem64** | -0.1 | 0.0851 | -0.267 | 0.0666 | 0.273 | 1 | loglinear | NM_181401.3:1125 |
| **Slco2b1** | 0.195 | 0.166 | -0.131 | 0.52 | 0.275 | 1 | lm.nb | NM_175316.3:2720 |
| **Rpl29** | -0.026 | 0.0223 | -0.0696 | 0.0176 | 0.277 | 1 | loglinear | NM_009082.2:110 |
| **Tet1** | 0.086 | 0.0738 | -0.0587 | 0.231 | 0.278 | 1 | loglinear | NM_027384.1:2192 |
| **Eif1** | 0.0403 | 0.0346 | -0.0276 | 0.108 | 0.278 | 1 | loglinear | NM_011508.1:664 |
| **Prkacb** | 0.0943 | 0.0811 | -0.0647 | 0.253 | 0.278 | 1 | loglinear | NM_011100.3:3754 |
| **Casp9** | 0.092 | 0.0791 | -0.0631 | 0.247 | 0.279 | 1 | loglinear | NM_015733.4:1675 |
| **Mre11a** | 0.107 | 0.0921 | -0.0736 | 0.287 | 0.279 | 1 | loglinear | NM_018736.2:2376 |
| **Timeless** | -0.129 | 0.112 | -0.348 | 0.0896 | 0.28 | 1 | loglinear | NM_011589.1:3720 |
| **Slc44a1** | 0.0899 | 0.0777 | -0.0624 | 0.242 | 0.281 | 1 | loglinear | NM_001159633.1:944 |
| **Cox5b** | 0.0503 | 0.0435 | -0.035 | 0.136 | 0.281 | 1 | loglinear | NM_009942.2:332 |
| **Bid** | 0.0973 | 0.0845 | -0.0684 | 0.263 | 0.283 | 1 | loglinear | NM_007544.3:1307 |
| **Cotl1** | -0.0578 | 0.0504 | -0.157 | 0.041 | 0.285 | 1 | loglinear | NM_028071.3:325 |
| **Ezh2** | -0.106 | 0.093 | -0.289 | 0.0761 | 0.286 | 1 | loglinear | NM_007971.2:425 |
| **Becn1** | -0.0375 | 0.0332 | -0.102 | 0.0275 | 0.291 | 1 | loglinear | NM_019584.3:1145 |
| **Hprt** | 0.126 | 0.111 | -0.0925 | 0.344 | 0.291 | 1 | loglinear | NM_013556.2:30 |
| **Csf1** | -0.0995 | 0.0882 | -0.272 | 0.0733 | 0.292 | 1 | loglinear | NM_001113530.1:833 |
| **Tspan18** | -0.0907 | 0.0809 | -0.249 | 0.0677 | 0.294 | 1 | loglinear | NM_183180.2:1008 |
| **Ppp3ca** | 0.0779 | 0.0698 | -0.0589 | 0.215 | 0.297 | 1 | loglinear | NM_008913.4:1675 |
| **Nlgn1** | 0.068 | 0.061 | -0.0516 | 0.188 | 0.298 | 1 | loglinear | NM_138666.3:1028 |
| **Igf1r** | -0.0589 | 0.0533 | -0.163 | 0.0455 | 0.301 | 1 | loglinear | NM_010513.2:3390 |
| **Trp73** | -0.169 | 0.154 | -0.471 | 0.133 | 0.304 | 1 | lm.nb | NM_011642.3:2632 |
| **Camk4** | 0.126 | 0.115 | -0.0988 | 0.351 | 0.304 | 1 | lm.nb | NM_009793.3:3280 |
| **Grm3** | 0.18 | 0.164 | -0.141 | 0.501 | 0.305 | 1 | lm.nb | NM_181850.2:2525 |
| **Hrk** | 0.137 | 0.125 | -0.108 | 0.381 | 0.305 | 1 | loglinear | NM_007545.2:3458 |
| **Nrp2** | -0.0416 | 0.0381 | -0.116 | 0.0331 | 0.307 | 1 | loglinear | NM_001077403.1:610 |
| **Slc6a1** | 0.116 | 0.106 | -0.093 | 0.324 | 0.309 | 1 | lm.nb | NM_178703.4:1865 |
| **Mmp14** | 0.109 | 0.101 | -0.0882 | 0.306 | 0.311 | 1 | loglinear | NM_008608.3:554 |

| | | | | | | | | |
|---|---|---|---|---|---|---|---|---|
| **Rps2** | -0.0298 | 0.0278 | -0.0843 | 0.0247 | 0.315 | 1 | loglinear | NM_008503.5:304 |
| **Kat2a** | 0.0952 | 0.089 | -0.0792 | 0.27 | 0.316 | 1 | loglinear | NM_020004.5:1700 |
| **Mcm6** | -0.121 | 0.114 | -0.344 | 0.102 | 0.318 | 1 | lm.nb | NM_008567.1:1118 |
| **Gadd45a** | 0.165 | 0.155 | -0.139 | 0.469 | 0.319 | 1 | loglinear | NM_007836.1:654 |
| **C1qb** | -0.104 | 0.0976 | -0.295 | 0.0877 | 0.319 | 1 | loglinear | NM_009777.2:865 |
| **Apc** | 0.0418 | 0.0397 | -0.036 | 0.12 | 0.323 | 1 | loglinear | NM_007462.3:645 |
| **Npl** | 0.131 | 0.125 | -0.113 | 0.375 | 0.325 | 1 | loglinear | NM_028749.1:600 |
| **Bnip3** | 0.106 | 0.102 | -0.0927 | 0.305 | 0.326 | 1 | loglinear | NM_009760.4:1108 |
| **Chst8** | -0.165 | 0.156 | -0.471 | 0.142 | 0.328 | 1 | Wald | NM_175140.4:1196 |
| **Wdr5** | 0.082 | 0.0789 | -0.0726 | 0.237 | 0.329 | 1 | loglinear | NM_080848.2:1704 |
| **Stmn1** | -0.0183 | 0.0177 | -0.0529 | 0.0163 | 0.331 | 1 | loglinear | NM_019641.3:595 |
| **Rad1** | -0.119 | 0.115 | -0.346 | 0.107 | 0.331 | 1 | loglinear | NM_011232.2:406 |
| **Kdm4a** | -0.0483 | 0.0469 | -0.14 | 0.0435 | 0.332 | 1 | loglinear | NM_172382.2:1675 |
| **Slc17a7** | -0.185 | 0.181 | -0.541 | 0.171 | 0.338 | 1 | loglinear | NM_182993.2:530 |
| **Vamp7** | 0.0432 | 0.0426 | -0.0403 | 0.127 | 0.341 | 1 | loglinear | NM_011515.4:390 |
| **Prkar2b** | 0.0539 | 0.0535 | -0.051 | 0.159 | 0.343 | 1 | loglinear | NM_011158.3:918 |
| **Pcna** | -0.0874 | 0.0869 | -0.258 | 0.0829 | 0.344 | 1 | loglinear | NM_011045.2:590 |
| **Ifitm2** | 0.106 | 0.106 | -0.101 | 0.314 | 0.344 | 1 | loglinear | NM_030694.1:87 |
| **Ldlrad3** | -0.0953 | 0.0948 | -0.281 | 0.0905 | 0.344 | 1 | loglinear | NM_178886.2:1570 |
| **Kmt2a** | -0.0445 | 0.0445 | -0.132 | 0.0427 | 0.346 | 1 | loglinear | NM_001081049.1:2080 |
| **Epg5** | 0.0966 | 0.0967 | -0.093 | 0.286 | 0.347 | 1 | loglinear | NM_001195633.1:2436 |
| **Fancg** | 0.1 | 0.101 | -0.0974 | 0.298 | 0.35 | 1 | loglinear | NM_053081.2:250 |
| **Mcm5** | -0.161 | 0.163 | -0.481 | 0.159 | 0.352 | 1 | lm.nb | NM_008566.2:2244 |
| **Bak1** | 0.0447 | 0.0455 | -0.0445 | 0.134 | 0.355 | 1 | loglinear | NM_007523.2:470 |
| **Mdm2** | -0.0632 | 0.0644 | -0.189 | 0.0631 | 0.356 | 1 | loglinear | NM_010786.4:1664 |
| **Pik3r1** | 0.0859 | 0.0878 | -0.0862 | 0.258 | 0.357 | 1 | loglinear | NM_001024955.1:5664 |
| **Jarid2** | -0.0688 | 0.0712 | -0.208 | 0.0708 | 0.362 | 1 | loglinear | NM_021878.2:2160 |
| **Eef2k** | 0.0844 | 0.0875 | -0.0871 | 0.256 | 0.363 | 1 | loglinear | NM_007908.3:1166 |
| **Tmem100** | 0.151 | 0.157 | -0.157 | 0.46 | 0.365 | 1 | loglinear | NM_026433.2:1350 |
| **Cdc7** | 0.0861 | 0.0898 | -0.0898 | 0.262 | 0.365 | 1 | loglinear | NM_001271566.1:2804 |
| **Nfkbie** | -0.228 | 0.236 | -0.689 | 0.234 | 0.366 | 1 | Wald | NM_008690.3:630 |
| **Prkdc** | -0.0665 | 0.07 | -0.204 | 0.0708 | 0.37 | 1 | loglinear | NM_011159.2:3108 |

| | | | | | | | | |
|---|---|---|---|---|---|---|---|---|
| **Smarca4** | -0.0487 | 0.0516 | -0.15 | 0.0524 | 0.372 | 1 | loglinear | NM_011417.2:3540 |
| **Cd68** | -0.135 | 0.143 | -0.415 | 0.145 | 0.376 | 1 | Wald | NM_009853.1:636 |
| **Lmnb1** | -0.0981 | 0.105 | -0.303 | 0.107 | 0.376 | 1 | loglinear | NM_010721.2:805 |
| **Hilpda** | 0.117 | 0.124 | -0.125 | 0.359 | 0.376 | 1 | Wald | NM_023516.5:236 |
| **Tm4sf1** | 0.198 | 0.211 | -0.216 | 0.612 | 0.377 | 1 | lm.nb | NM_008536.3:652 |
| **Tcirg1** | 0.146 | 0.155 | -0.158 | 0.451 | 0.378 | 1 | Wald | NM_001136091.1:1345 |
| **Igf2r** | -0.0455 | 0.0488 | -0.141 | 0.0501 | 0.378 | 1 | loglinear | NM_010515.1:2585 |
| **Snca** | 0.123 | 0.133 | -0.138 | 0.384 | 0.384 | 1 | lm.nb | NM_009221.2:285 |
| **Rpl9** | -0.0285 | 0.0313 | -0.0898 | 0.0328 | 0.389 | 1 | loglinear | NM_011292.2:142 |
| **Ppp3r1** | 0.0335 | 0.0372 | -0.0394 | 0.106 | 0.394 | 1 | loglinear | NM_024459.2:1390 |
| **Traf6** | 0.0832 | 0.0926 | -0.0983 | 0.265 | 0.395 | 1 | loglinear | NM_009424.2:980 |
| **Irak1** | 0.0966 | 0.108 | -0.115 | 0.308 | 0.397 | 1 | loglinear | NM_008363.2:951 |
| **Ets2** | 0.0964 | 0.108 | -0.115 | 0.308 | 0.398 | 1 | loglinear | NM_011809.2:3284 |
| **Vim** | -0.0898 | 0.101 | -0.287 | 0.107 | 0.398 | 1 | loglinear | NM_011701.4:34 |
| **Ncor1** | 0.043 | 0.0485 | -0.052 | 0.138 | 0.401 | 1 | loglinear | NM_011308.2:211 |
| **Ptx3** | -0.0943 | 0.106 | -0.303 | 0.114 | 0.401 | 1 | loglinear | NM_008987.3:692 |
| **Nfkbia** | -0.146 | 0.163 | -0.466 | 0.175 | 0.402 | 1 | Wald | NM_010907.2:646 |
| **Ambra1** | -0.0284 | 0.0321 | -0.0913 | 0.0346 | 0.403 | 1 | loglinear | NM_001080754.1:944 |
| **Prkce** | -0.0874 | 0.0991 | -0.282 | 0.107 | 0.404 | 1 | loglinear | NM_011104.2:1510 |
| **Cd109** | 0.147 | 0.165 | -0.177 | 0.47 | 0.404 | 1 | Wald | NM_153098.3:2720 |
| **Kdm1a** | -0.0278 | 0.0316 | -0.0898 | 0.0342 | 0.405 | 1 | loglinear | NM_133872.1:1263 |
| **Maff** | -0.156 | 0.176 | -0.502 | 0.19 | 0.406 | 1 | Wald | NM_010755.3:743 |
| **Flt1** | 0.131 | 0.149 | -0.16 | 0.422 | 0.407 | 1 | Wald | NM_010228.3:1550 |
| **Brd4** | -0.0379 | 0.0436 | -0.123 | 0.0475 | 0.41 | 1 | loglinear | NM_001286630.1:1492 |
| **Ezh1** | 0.148 | 0.17 | -0.185 | 0.481 | 0.41 | 1 | loglinear | NM_007970.1:3950 |
| **Dnmt1** | 0.074 | 0.0854 | -0.0935 | 0.241 | 0.412 | 1 | loglinear | NM_010066.3:2380 |
| **Trem2** | -0.202 | 0.233 | -0.658 | 0.254 | 0.414 | 1 | Wald | NM_031254.2:646 |
| **Optn** | 0.094 | 0.11 | -0.121 | 0.309 | 0.417 | 1 | loglinear | NM_181848.4:1018 |
| **Eed** | -0.0356 | 0.0417 | -0.117 | 0.0461 | 0.418 | 1 | loglinear | NM_021876.3:1000 |
| **Calr** | -0.0479 | 0.0567 | -0.159 | 0.0633 | 0.423 | 1 | loglinear | NM_007591.3:551 |
| **Fscn1** | -0.022 | 0.0261 | -0.073 | 0.0291 | 0.424 | 1 | loglinear | NM_007984.2:1645 |
| **Mbd3** | 0.0574 | 0.0685 | -0.0768 | 0.192 | 0.426 | 1 | loglinear | NM_013595.2:420 |

| | | | | | | | | |
|---|---|---|---|---|---|---|---|---|
| **Ugt8a** | 0.129 | 0.153 | -0.17 | 0.428 | 0.426 | 1 | Wald | NM_011674.4:138 |
| **Dab2** | -0.128 | 0.152 | -0.426 | 0.17 | 0.427 | 1 | Wald | NM_023118.2:415 |
| **Tmcc3** | 0.113 | 0.135 | -0.152 | 0.378 | 0.427 | 1 | loglinear | NM_172051.2:3825 |
| **Homer1** | 0.0539 | 0.0647 | -0.0729 | 0.181 | 0.429 | 1 | loglinear | NM_147176.2:1165 |
| **Tgfa** | 0.0922 | 0.111 | -0.126 | 0.31 | 0.431 | 1 | loglinear | NM_031199.2:3360 |
| **Pdpn** | -0.0782 | 0.0946 | -0.264 | 0.107 | 0.432 | 1 | loglinear | NM_010329.2:1625 |
| **Chek2** | 0.124 | 0.149 | -0.169 | 0.416 | 0.435 | 1 | Wald | NM_016681.3:790 |
| **Suv39h2** | -0.0834 | 0.101 | -0.282 | 0.115 | 0.435 | 1 | loglinear | NM_022724.4:1427 |
| **Cnn2** | -0.124 | 0.151 | -0.419 | 0.172 | 0.436 | 1 | loglinear | NM_007725.2:350 |
| **Igsf10** | 0.135 | 0.165 | -0.189 | 0.459 | 0.438 | 1 | lm.nb | NM_001162884.1:3100 |
| **Kdm5c** | -0.0976 | 0.12 | -0.333 | 0.138 | 0.44 | 1 | lm.nb | NM_013668.3:750 |
| **Kif2c** | -0.175 | 0.217 | -0.6 | 0.249 | 0.442 | 1 | lm.nb | NM_134471.3:915 |
| **Pole** | -0.0746 | 0.0922 | -0.255 | 0.106 | 0.442 | 1 | loglinear | NM_011132.2:823 |
| **Lingo1** | 0.0651 | 0.0814 | -0.0944 | 0.225 | 0.447 | 1 | loglinear | NM_181074.4:1087 |
| **Ctsw** | -0.17 | 0.214 | -0.589 | 0.249 | 0.453 | 1 | Wald | NM_009985.4:190 |
| **Ikbkb** | -0.0362 | 0.0459 | -0.126 | 0.0539 | 0.454 | 1 | loglinear | NM_010546.2:498 |
| **Clstn1** | 0.0568 | 0.0724 | -0.0852 | 0.199 | 0.456 | 1 | loglinear | NM_023051.4:2352 |
| **Srxn1** | 0.106 | 0.135 | -0.159 | 0.371 | 0.457 | 1 | loglinear | NM_029688.4:2010 |
| **Kdm1b** | 0.0574 | 0.0738 | -0.0872 | 0.202 | 0.459 | 1 | loglinear | NM_172262.3:2034 |
| **Emp1** | -0.0973 | 0.127 | -0.346 | 0.151 | 0.464 | 1 | loglinear | NM_010128.4:1080 |
| **Stx18** | -0.135 | 0.177 | -0.481 | 0.211 | 0.467 | 1 | lm.nb | NM_026959.2:770 |
| **Plekhm1** | -0.0445 | 0.0584 | -0.159 | 0.0699 | 0.468 | 1 | loglinear | NM_183034.1:3060 |
| **Kdm4b** | -0.0321 | 0.0425 | -0.115 | 0.0512 | 0.472 | 1 | loglinear | NM_172132.1:3704 |
| **Nrgn** | 0.216 | 0.29 | -0.353 | 0.784 | 0.479 | 1 | lm.nb | NM_022029.2:192 |
| **Esam** | -0.0557 | 0.0752 | -0.203 | 0.0916 | 0.48 | 1 | loglinear | NM_027102.3:495 |
| **Tarbp2** | 0.0623 | 0.0843 | -0.103 | 0.227 | 0.481 | 1 | loglinear | NM_001253795.1:813 |
| **Rab7** | 0.0262 | 0.0354 | -0.0433 | 0.0956 | 0.481 | 1 | loglinear | NM_009005.2:490 |
| **Grm2** | -0.0698 | 0.0946 | -0.255 | 0.116 | 0.481 | 1 | loglinear | NM_001160353.1:2770 |
| **Kdm5a** | -0.0642 | 0.0872 | -0.235 | 0.107 | 0.482 | 1 | loglinear | XR_377436.1:2314 |
| **Asph** | 0.0507 | 0.0692 | -0.0848 | 0.186 | 0.484 | 1 | loglinear | NM_001177849.1:400 |
| **Chek1** | 0.0983 | 0.134 | -0.164 | 0.36 | 0.486 | 1 | Wald | NM_007691.5:1220 |
| **Tubb4a** | -0.0838 | 0.116 | -0.312 | 0.144 | 0.492 | 1 | loglinear | NM_009451.3:1819 |

| | | | | | | | | |
|---|---|---|---|---|---|---|---|---|
| **Bad** | -0.0351 | 0.0488 | -0.131 | 0.0606 | 0.493 | 1 | loglinear | NM_007522.3:1146 |
| **Lrrc3** | 0.0846 | 0.118 | -0.148 | 0.317 | 0.495 | 1 | lm.nb | NM_145152.4:1830 |
| **Col6a3** | -0.166 | 0.236 | -0.628 | 0.296 | 0.501 | 1 | lm.nb | XM_897036.2:4583 |
| **Setd7** | -0.078 | 0.111 | -0.295 | 0.139 | 0.501 | 1 | loglinear | NM_080793.5:3905 |
| **Txnrd1** | -0.0344 | 0.0491 | -0.131 | 0.0618 | 0.503 | 1 | loglinear | NM_015762.2:2245 |
| **Cldn5** | 0.0604 | 0.086 | -0.108 | 0.229 | 0.503 | 1 | loglinear | NM_013805.4:975 |
| **Ddx58** | -0.108 | 0.153 | -0.408 | 0.193 | 0.505 | 1 | Wald | NM_172689.3:1751 |
| **Pms2** | 0.0508 | 0.073 | -0.0923 | 0.194 | 0.506 | 1 | loglinear | NM_008886.2:265 |
| **Shank3** | -0.103 | 0.148 | -0.394 | 0.188 | 0.508 | 1 | lm.nb | NM_021423.3:1855 |
| **Cnp** | -0.0814 | 0.118 | -0.312 | 0.149 | 0.509 | 1 | lm.nb | NM_009923.2:166 |
| **Brd3** | -0.035 | 0.0507 | -0.134 | 0.0643 | 0.509 | 1 | loglinear | NM_001113573.1:2690 |
| **Ercc2** | 0.0351 | 0.051 | -0.0648 | 0.135 | 0.51 | 1 | loglinear | NM_007949.4:1800 |
| **Lamp2** | -0.0368 | 0.0535 | -0.142 | 0.068 | 0.511 | 1 | loglinear | NM_001017959.1:908 |
| **Mafb** | -0.0719 | 0.105 | -0.278 | 0.134 | 0.512 | 1 | loglinear | NM_010658.2:2658 |
| **Jam2** | 0.04 | 0.0584 | -0.0745 | 0.155 | 0.513 | 1 | loglinear | NM_023844.4:1090 |
| **Dnmt3a** | -0.0669 | 0.0986 | -0.26 | 0.126 | 0.516 | 1 | loglinear | NM_007872.4:7160 |
| **Rala** | 0.0297 | 0.0443 | -0.0572 | 0.117 | 0.522 | 1 | loglinear | NM_019491.5:605 |
| **Fancc** | -0.0574 | 0.0865 | -0.227 | 0.112 | 0.526 | 1 | loglinear | NM_007985.2:1115 |
| **Cyp7b1** | -0.107 | 0.163 | -0.427 | 0.212 | 0.531 | 1 | Wald | NM_007825.4:1030 |
| **Gstm1** | -0.117 | 0.179 | -0.468 | 0.233 | 0.533 | 1 | Wald | NM_010358.5:50 |
| **Icam2** | -0.103 | 0.158 | -0.412 | 0.207 | 0.533 | 1 | loglinear | NM_010494.1:375 |
| **Trpm4** | 0.0802 | 0.123 | -0.161 | 0.322 | 0.536 | 1 | Wald | NM_175130.4:1145 |
| **Bcl10** | -0.0562 | 0.0869 | -0.227 | 0.114 | 0.536 | 1 | loglinear | NM_009740.1:1168 |
| **Kdm6a** | -0.111 | 0.172 | -0.448 | 0.227 | 0.538 | 1 | lm.nb | NM_009483.1:2560 |
| **Bin1** | -0.0191 | 0.0301 | -0.0781 | 0.0398 | 0.542 | 1 | loglinear | NM_001083334.1:1400 |
| **Bok** | 0.0594 | 0.0935 | -0.124 | 0.243 | 0.543 | 1 | loglinear | NM_016778.2:635 |
| **Sod2** | -0.0445 | 0.0702 | -0.182 | 0.093 | 0.543 | 1 | loglinear | NM_013671.3:1495 |
| **Bcl2l1** | -0.0402 | 0.0634 | -0.164 | 0.084 | 0.543 | 1 | loglinear | NM_009743.4:200 |
| **Pttg1** | 0.0826 | 0.13 | -0.173 | 0.338 | 0.545 | 1 | loglinear | NM_001131054.1:288 |
| **Csf1r** | -0.0903 | 0.144 | -0.372 | 0.192 | 0.548 | 1 | loglinear | NM_001037859.1:3655 |
| **Ngfr** | 0.0792 | 0.127 | -0.169 | 0.327 | 0.549 | 1 | loglinear | NM_033217.3:1995 |
| **Ripk2** | 0.0759 | 0.123 | -0.165 | 0.317 | 0.554 | 1 | loglinear | NM_138952.3:830 |

| | | | | | | | | |
|---|---|---|---|---|---|---|---|---|
| **Bcl2** | -0.0622 | 0.101 | -0.26 | 0.135 | 0.554 | 1 | loglinear | NM_009741.3:1844 |
| **Serpine1** | -0.145 | 0.235 | -0.605 | 0.315 | 0.555 | 1 | Wald | NM_008871.2:1822 |
| **Gria1** | -0.0376 | 0.0614 | -0.158 | 0.0826 | 0.557 | 1 | loglinear | NM_001252403.1:2476 |
| **Dicer1** | 0.0341 | 0.0558 | -0.0752 | 0.143 | 0.558 | 1 | loglinear | NM_148948.2:5270 |
| **Spp1** | -0.119 | 0.197 | -0.505 | 0.267 | 0.563 | 1 | lm.nb | NM_009263.3:420 |
| **St3gal6** | 0.0958 | 0.159 | -0.215 | 0.407 | 0.563 | 1 | loglinear | NM_018784.2:1252 |
| **Gsn** | -0.105 | 0.174 | -0.445 | 0.236 | 0.564 | 1 | lm.nb | NM_146120.3:624 |
| **Ifitm3** | -0.0948 | 0.158 | -0.405 | 0.215 | 0.565 | 1 | lm.nb | NM_025378.2:370 |
| **Chuk** | -0.0227 | 0.0381 | -0.0973 | 0.0519 | 0.567 | 1 | loglinear | NM_001162410.1:222 |
| **Cables1** | 0.0409 | 0.0693 | -0.0949 | 0.177 | 0.572 | 1 | loglinear | NM_001146287.1:3120 |
| **Sox4** | -0.0425 | 0.0723 | -0.184 | 0.0992 | 0.573 | 1 | loglinear | NM_009238.2:2635 |
| **F3** | 0.0541 | 0.0921 | -0.126 | 0.235 | 0.573 | 1 | loglinear | NM_010171.3:1170 |
| **Sin3a** | -0.0502 | 0.0858 | -0.218 | 0.118 | 0.574 | 1 | loglinear | NM_001110350.1:3585 |
| **Amigo2** | 0.0517 | 0.0883 | -0.121 | 0.225 | 0.574 | 1 | loglinear | NM_178114.4:2425 |
| **Smarcd1** | -0.0676 | 0.116 | -0.295 | 0.16 | 0.576 | 1 | loglinear | NM_031842.1:2220 |
| **Msh2** | 0.0373 | 0.0645 | -0.0891 | 0.164 | 0.579 | 1 | loglinear | NM_008628.2:1870 |
| **Pik3r2** | 0.029 | 0.0502 | -0.0694 | 0.127 | 0.579 | 1 | loglinear | NM_008841.2:230 |
| **Map2k1** | -0.0571 | 0.0995 | -0.252 | 0.138 | 0.582 | 1 | loglinear | NM_008927.3:1695 |
| **E2f1** | -0.0839 | 0.146 | -0.371 | 0.203 | 0.582 | 1 | lm.nb | NM_007891.4:926 |
| **Mal** | -0.136 | 0.239 | -0.605 | 0.333 | 0.587 | 1 | Wald | NM_001171187.1:685 |
| **B3gnt5** | -0.0338 | 0.0601 | -0.152 | 0.0839 | 0.589 | 1 | loglinear | NM_001159407.1:1738 |
| **Hira** | -0.0299 | 0.0532 | -0.134 | 0.0743 | 0.589 | 1 | loglinear | NM_010435.2:3210 |
| **Map2k4** | 0.0341 | 0.0609 | -0.0852 | 0.153 | 0.591 | 1 | loglinear | NM_009157.4:1335 |
| **Abl1** | -0.0431 | 0.077 | -0.194 | 0.108 | 0.591 | 1 | loglinear | NM_009594.4:1378 |
| **Iqsec1** | 0.0278 | 0.0505 | -0.0712 | 0.127 | 0.597 | 1 | loglinear | NM_001134383.1:1792 |
| **Atg5** | 0.0354 | 0.0647 | -0.0914 | 0.162 | 0.599 | 1 | loglinear | NM_001314013.1:262 |
| **Rpl36al** | -0.0196 | 0.036 | -0.0901 | 0.0509 | 0.601 | 1 | loglinear | NM_025589.4:74 |
| **Itgav** | 0.0498 | 0.0921 | -0.131 | 0.23 | 0.603 | 1 | loglinear | NM_008402.2:3145 |
| **Grin2b** | 0.0468 | 0.0878 | -0.125 | 0.219 | 0.608 | 1 | loglinear | NM_008171.3:6340 |
| **Rapgef3** | -0.072 | 0.136 | -0.338 | 0.194 | 0.61 | 1 | lm.nb | NM_001177810.1:564 |
| **Atg7** | 0.0562 | 0.106 | -0.152 | 0.265 | 0.612 | 1 | loglinear | NM_028835.1:855 |
| **Mr1** | 0.105 | 0.198 | -0.284 | 0.494 | 0.613 | 1 | Wald | NM_008209.4:1360 |

| | | | | | | | | |
|---|---|---|---|---|---|---|---|---|
| **Irf8** | -0.12 | 0.227 | -0.564 | 0.325 | 0.614 | 1 | Wald | NM_008320.3:2274 |
| **Kdm4c** | -0.0244 | 0.0466 | -0.116 | 0.067 | 0.615 | 1 | loglinear | NM_144787.1:2859 |
| **Ak1** | 0.0311 | 0.0602 | -0.0869 | 0.149 | 0.619 | 1 | loglinear | NM_001198790.1:1625 |
| **Cdkn1c** | -0.0589 | 0.114 | -0.283 | 0.165 | 0.621 | 1 | lm.nb | NM_009876.3:1240 |
| **Rrm2** | -0.083 | 0.162 | -0.4 | 0.234 | 0.622 | 1 | lm.nb | NM_009104.1:265 |
| **Pld2** | -0.0684 | 0.133 | -0.329 | 0.193 | 0.624 | 1 | Wald | NM_008876.2:1134 |
| **Rad9a** | -0.0852 | 0.167 | -0.413 | 0.242 | 0.626 | 1 | Wald | NM_011237.2:490 |
| **Parp1** | -0.0246 | 0.0489 | -0.121 | 0.0713 | 0.629 | 1 | loglinear | NM_007415.2:3020 |
| **Ikbke** | 0.0985 | 0.196 | -0.285 | 0.482 | 0.63 | 1 | Wald | NM_019777.3:618 |
| **Birc5** | -0.0736 | 0.148 | -0.363 | 0.216 | 0.631 | 1 | loglinear | NM_009689.2:237 |
| **Prkaca** | 0.0179 | 0.0368 | -0.0542 | 0.0901 | 0.639 | 1 | loglinear | NM_008854.3:699 |
| **Fcgr1** | 0.0906 | 0.188 | -0.278 | 0.459 | 0.644 | 1 | Wald | NM_010186.5:185 |
| **Cln3** | 0.0638 | 0.134 | -0.199 | 0.326 | 0.646 | 1 | lm.nb | NM_001146311.1:378 |
| **Apoe** | 0.115 | 0.245 | -0.364 | 0.595 | 0.65 | 1 | lm.nb | NM_001305844.1:903 |
| **Irf1** | -0.0926 | 0.198 | -0.481 | 0.295 | 0.654 | 1 | Wald | NM_008390.1:365 |
| **Mfge8** | 0.0625 | 0.138 | -0.207 | 0.332 | 0.662 | 1 | loglinear | NM_008594.2:1357 |
| **Dot1l** | -0.037 | 0.082 | -0.198 | 0.124 | 0.664 | 1 | loglinear | NM_199322.1:5490 |
| **Pld1** | 0.059 | 0.13 | -0.197 | 0.315 | 0.665 | 1 | Wald | NM_001164056.1:520 |
| **Crebbp** | 0.0169 | 0.0377 | -0.0569 | 0.0908 | 0.665 | 1 | loglinear | NM_001025432.1:3770 |
| **Atr** | -0.0355 | 0.0791 | -0.191 | 0.12 | 0.666 | 1 | loglinear | NM_019864.1:4392 |
| **Gria2** | 0.0189 | 0.0424 | -0.0642 | 0.102 | 0.668 | 1 | loglinear | NM_001039195.1:300 |
| **S100a10** | -0.0425 | 0.0954 | -0.229 | 0.144 | 0.668 | 1 | loglinear | NM_009112.2:154 |
| **Gja1** | -0.0864 | 0.195 | -0.468 | 0.296 | 0.669 | 1 | lm.nb | NM_010288.3:1450 |
| **Ep300** | -0.0167 | 0.0379 | -0.0909 | 0.0575 | 0.671 | 1 | loglinear | NM_177821.6:4305 |
| **Tpd52** | 0.0299 | 0.0695 | -0.106 | 0.166 | 0.678 | 1 | loglinear | NM_001025262.1:2054 |
| **Nthl1** | -0.0593 | 0.137 | -0.328 | 0.209 | 0.678 | 1 | Wald | NM_008743.2:34 |
| **Emcn** | -0.0405 | 0.0974 | -0.232 | 0.15 | 0.688 | 1 | loglinear | NM_001163522.1:230 |
| **Sox9** | -0.0268 | 0.0648 | -0.154 | 0.1 | 0.69 | 1 | loglinear | NM_011448.4:3540 |
| **Setd1b** | -0.028 | 0.068 | -0.161 | 0.105 | 0.691 | 1 | loglinear | NM_001040398.1:2880 |
| **Cflar** | 0.0447 | 0.109 | -0.169 | 0.259 | 0.693 | 1 | loglinear | NM_207653.3:990 |
| **Ccng2** | -0.0293 | 0.0715 | -0.169 | 0.111 | 0.693 | 1 | loglinear | NM_007635.4:1536 |
| **Cp** | 0.0566 | 0.143 | -0.223 | 0.336 | 0.702 | 1 | loglinear | NM_001042611.1:1750 |

| | | | | | | | | |
|---|---|---|---|---|---|---|---|---|
| **Sftpd** | -0.0697 | 0.176 | -0.415 | 0.276 | 0.704 | 1 | Wald | NM_009160.2:85 |
| **Ago4** | 0.0398 | 0.102 | -0.16 | 0.239 | 0.706 | 1 | loglinear | NM_153177.3:1576 |
| **Atg9a** | -0.0219 | 0.0562 | -0.132 | 0.0882 | 0.707 | 1 | loglinear | NM_001003917.3:340 |
| **Atp6v1a** | 0.0311 | 0.0814 | -0.129 | 0.191 | 0.713 | 1 | loglinear | NM_007508.5:434 |
| **Blm** | -0.0435 | 0.116 | -0.271 | 0.184 | 0.717 | 1 | loglinear | NM_001042527.2:264 |
| **Gdpd2** | -0.0283 | 0.0768 | -0.179 | 0.122 | 0.722 | 1 | loglinear | NM_023608.3:1438 |
| **Map3k1** | 0.0255 | 0.0698 | -0.111 | 0.162 | 0.724 | 1 | loglinear | NM_011945.2:1640 |
| **Nfkb1** | 0.023 | 0.0645 | -0.103 | 0.149 | 0.73 | 1 | loglinear | NM_008689.2:2125 |
| **Gadd45g** | -0.0676 | 0.19 | -0.439 | 0.304 | 0.731 | 1 | lm.nb | NM_011817.2:208 |
| **Mbd2** | -0.013 | 0.0368 | -0.0851 | 0.0592 | 0.733 | 1 | loglinear | NM_010773.2:655 |
| **Sqstm1** | 0.0241 | 0.0687 | -0.111 | 0.159 | 0.735 | 1 | loglinear | NM_011018.2:1430 |
| **Psen2** | 0.0453 | 0.129 | -0.207 | 0.298 | 0.736 | 1 | Wald | NM_001128605.1:560 |
| **Rtn4rl1** | -0.063 | 0.18 | -0.417 | 0.291 | 0.736 | 1 | lm.nb | NM_177708.4:2075 |
| **Tbc1d4** | 0.0434 | 0.124 | -0.2 | 0.287 | 0.737 | 1 | Wald | NM_001081278.2:2610 |
| **Gba** | 0.0514 | 0.152 | -0.246 | 0.349 | 0.743 | 1 | loglinear | NM_001077411.1:820 |
| **Ncaph** | -0.0454 | 0.134 | -0.308 | 0.217 | 0.744 | 1 | lm.nb | NM_144818.3:1540 |
| **Fadd** | -0.0475 | 0.141 | -0.325 | 0.23 | 0.747 | 1 | Wald | NM_010175.5:2641 |
| **Nefl** | 0.0569 | 0.171 | -0.277 | 0.391 | 0.747 | 1 | lm.nb | NM_010910.1:1303 |
| **Tnfrsf25** | 0.0558 | 0.167 | -0.271 | 0.382 | 0.747 | 1 | Wald | NM_033042.3:685 |
| **Irf3** | -0.0256 | 0.078 | -0.178 | 0.127 | 0.751 | 1 | loglinear | NM_016849.4:526 |
| **Mvp** | -0.0599 | 0.182 | -0.416 | 0.296 | 0.751 | 1 | Wald | NM_080638.2:1845 |
| **Atg14** | 0.0312 | 0.0956 | -0.156 | 0.219 | 0.752 | 1 | loglinear | NM_172599.4:204 |
| **Clic4** | -0.0337 | 0.104 | -0.237 | 0.169 | 0.754 | 1 | loglinear | NM_013885.2:3280 |
| **Sumo1** | 0.0133 | 0.0415 | -0.0681 | 0.0947 | 0.757 | 1 | loglinear | NM_009460.1:720 |
| **Ltbr** | -0.0404 | 0.125 | -0.286 | 0.205 | 0.757 | 1 | Wald | NM_010736.3:1962 |
| **Mef2c** | -0.0439 | 0.142 | -0.323 | 0.235 | 0.766 | 1 | lm.nb | NM_001170537.1:4341 |
| **Fyn** | 0.00693 | 0.0232 | -0.0386 | 0.0524 | 0.773 | 1 | loglinear | NM_008054.2:1030 |
| **C3** | 0.071 | 0.24 | -0.399 | 0.541 | 0.776 | 1 | Wald | XM_011246258.1:2702 |
| **Lyn** | 0.0375 | 0.128 | -0.214 | 0.289 | 0.778 | 1 | Wald | NM_010747.1:1725 |
| **Sirt1** | 0.0203 | 0.0729 | -0.123 | 0.163 | 0.788 | 1 | loglinear | NM_019812.2:843 |
| **Pacsin1** | 0.0341 | 0.126 | -0.214 | 0.282 | 0.794 | 1 | lm.nb | NM_011861.3:2936 |
| **Grap** | -0.0458 | 0.169 | -0.378 | 0.286 | 0.795 | 1 | Wald | NM_027817.3:121 |

| | | | | | | | | |
|---|---|---|---|---|---|---|---|---|
| **Lig1** | -0.0257 | 0.0971 | -0.216 | 0.165 | 0.798 | 1 | loglinear | NM_001083188.1:1456 |
| **Anapc15** | -0.0275 | 0.104 | -0.232 | 0.177 | 0.799 | 1 | loglinear | NM_027532.3:350 |
| **Gclc** | 0.0294 | 0.113 | -0.191 | 0.25 | 0.801 | 1 | loglinear | NM_010295.2:1102 |
| **Kdm2b** | -0.00866 | 0.0339 | -0.075 | 0.0577 | 0.805 | 1 | loglinear | NM_001003953.1:3984 |
| **Kcnk13** | -0.418 | 1.64 | -3.64 | 2.8 | 0.807 | 1 | Wald | NM_146037.1:1430 |
| **Uty** | 0.642 | 2.56 | -4.38 | 5.67 | 0.809 | 1 | loglinear | NM_009484.2:3530 |
| **Dlg4** | 0.0171 | 0.0688 | -0.118 | 0.152 | 0.81 | 1 | loglinear | NM_001109752.1:1866 |
| **Bcl2l2** | -0.0109 | 0.0442 | -0.0975 | 0.0757 | 0.811 | 1 | loglinear | NM_007537.1:1592 |
| **Nwd1** | 0.0216 | 0.0889 | -0.153 | 0.196 | 0.814 | 1 | loglinear | NM_176940.5:3564 |
| **Cd24a** | -0.0109 | 0.0453 | -0.0996 | 0.0779 | 0.817 | 1 | loglinear | NM_009846.2:584 |
| **Ulk1** | -0.0113 | 0.048 | -0.105 | 0.0828 | 0.819 | 1 | loglinear | NM_009469.3:4050 |
| **Kdm5d** | 0.627 | 2.67 | -4.6 | 5.85 | 0.82 | 1 | loglinear | NM_011419.3:2390 |
| **Map3k14** | -0.0473 | 0.2 | -0.44 | 0.345 | 0.82 | 1 | Wald | NM_016896.3:3930 |
| **St8sia6** | 0.0365 | 0.155 | -0.268 | 0.341 | 0.821 | 1 | Wald | NM_145838.1:1578 |
| **Osmr** | -0.0364 | 0.158 | -0.346 | 0.273 | 0.824 | 1 | Wald | NM_011019.3:395 |
| **Sesn2** | -0.0457 | 0.201 | -0.439 | 0.348 | 0.827 | 1 | Wald | NM_144907.1:1416 |
| **Lfng** | 0.00862 | 0.0384 | -0.0667 | 0.0839 | 0.828 | 1 | loglinear | NM_008494.3:1965 |
| **Nlgn2** | -0.00657 | 0.0294 | -0.0642 | 0.0511 | 0.829 | 1 | loglinear | NM_198862.2:1204 |
| **Kdm3a** | -0.0205 | 0.0934 | -0.204 | 0.162 | 0.832 | 1 | loglinear | NM_001038695.2:4110 |
| **Lilrb4a** | -0.536 | 2.45 | -5.34 | 4.27 | 0.833 | 1 | Wald | NM_013532.3:348 |
| **Map1lc3a** | 0.0178 | 0.0867 | -0.152 | 0.188 | 0.843 | 1 | loglinear | NM_025735.1:685 |
| **Rb1cc1** | 0.00562 | 0.0288 | -0.0509 | 0.0621 | 0.85 | 1 | loglinear | NM_009826.4:497 |
| **Mpeg1** | 0.0275 | 0.141 | -0.249 | 0.304 | 0.851 | 1 | Wald | NM_010821.1:4135 |
| **Ngf** | -0.474 | 2.44 | -5.25 | 4.3 | 0.851 | 1 | Wald | NM_001112698.1:630 |
| **Irf2** | 0.0269 | 0.142 | -0.25 | 0.304 | 0.854 | 1 | lm.nb | NM_008391.2:440 |
| **Dusp7** | 0.0188 | 0.1 | -0.177 | 0.215 | 0.855 | 1 | loglinear | NM_153459.4:2094 |
| **Ldha** | -0.0173 | 0.0945 | -0.203 | 0.168 | 0.859 | 1 | loglinear | NM_010699.2:1354 |
| **Ifi30** | -0.0184 | 0.101 | -0.216 | 0.18 | 0.86 | 1 | loglinear | NM_023065.3:806 |
| **Grn** | -0.0141 | 0.079 | -0.169 | 0.141 | 0.862 | 1 | loglinear | NM_008175.3:2010 |
| **Trp53** | 0.0199 | 0.115 | -0.206 | 0.246 | 0.867 | 1 | loglinear | NM_011640.1:1835 |
| **Mapt** | -0.013 | 0.0792 | -0.168 | 0.142 | 0.873 | 1 | loglinear | NM_001038609.2:1202 |
| **Ogg1** | -0.0248 | 0.15 | -0.319 | 0.27 | 0.874 | 1 | Wald | NM_010957.4:168 |

| | | | | | | | | |
|---|---|---|---|---|---|---|---|---|
| **Rnf8** | 0.0072 | 0.0457 | -0.0824 | 0.0968 | 0.879 | 1 | loglinear | NM_021419.2:1671 |
| **Reln** | -0.00788 | 0.053 | -0.112 | 0.0961 | 0.886 | 1 | loglinear | NM_011261.2:2545 |
| **Prdx1** | 0.0263 | 0.177 | -0.321 | 0.374 | 0.886 | 1 | lm.nb | NM_011034.4:1131 |
| **Rpa1** | -0.00706 | 0.0479 | -0.101 | 0.0867 | 0.886 | 1 | loglinear | NM_026653.2:930 |
| **Rab6b** | 0.0137 | 0.0937 | -0.17 | 0.197 | 0.887 | 1 | loglinear | NM_173781.4:715 |
| **Tnfsf12** | 0.331 | 2.32 | -4.22 | 4.88 | 0.891 | 1 | Wald | NM_011614.3:1215 |
| **Pten** | 0.00795 | 0.0561 | -0.102 | 0.118 | 0.891 | 1 | loglinear | NM_008960.2:5160 |
| **Birc3** | 0.0334 | 0.244 | -0.445 | 0.511 | 0.895 | 1 | Wald | NM_007464.3:425 |
| **Cd163** | -0.44 | 3.22 | -6.76 | 5.88 | 0.895 | 1 | Wald | NM_053094.2:3225 |
| **Tmem88b** | 0.0238 | 0.184 | -0.336 | 0.384 | 0.9 | 1 | Wald | NM_001033394.3:1122 |
| **Gpr183** | -0.345 | 2.78 | -5.8 | 5.11 | 0.905 | 1 | Wald | NM_183031.2:238 |
| **Cks1b** | -0.015 | 0.127 | -0.263 | 0.233 | 0.909 | 1 | loglinear | NM_016904.1:185 |
| **Ptpn6** | 0.0255 | 0.216 | -0.399 | 0.45 | 0.909 | 1 | Wald | NM_013545.2:1691 |
| **Cd84** | -0.458 | 3.93 | -8.16 | 7.24 | 0.91 | 1 | Wald | NM_013489.2:915 |
| **Ctsf** | 0.0128 | 0.112 | -0.206 | 0.232 | 0.912 | 1 | loglinear | NM_019861.1:625 |
| **C1qc** | -0.0135 | 0.119 | -0.246 | 0.219 | 0.912 | 1 | loglinear | NM_007574.2:708 |
| **Msr1** | -0.304 | 2.67 | -5.53 | 4.92 | 0.912 | 1 | Wald | NM_001113326.1:555 |
| **Tgfbr1** | -0.00644 | 0.0585 | -0.121 | 0.108 | 0.915 | 1 | loglinear | NM_009370.2:4425 |
| **Setdb1** | -0.0091 | 0.0851 | -0.176 | 0.158 | 0.917 | 1 | loglinear | NM_018877.2:1625 |
| **Pik3cb** | 0.00527 | 0.0492 | -0.0912 | 0.102 | 0.917 | 1 | loglinear | NM_029094.3:1970 |
| **Kir3dl2** | 0.0252 | 0.235 | -0.435 | 0.486 | 0.918 | 1 | Wald | NM_177748.2:1519 |
| **Ccr5** | 0.315 | 2.94 | -5.46 | 6.09 | 0.918 | 1 | Wald | NM_009917.5:1340 |
| **Bag4** | 0.00945 | 0.0905 | -0.168 | 0.187 | 0.919 | 1 | loglinear | NM_026121.3:3735 |
| **Setd2** | -0.00256 | 0.0249 | -0.0513 | 0.0462 | 0.921 | 1 | loglinear | NM_001081340.2:1345 |
| **Pik3cd** | -0.00614 | 0.0606 | -0.125 | 0.113 | 0.922 | 1 | loglinear | XM_003945690.1:4648 |
| **Grin2a** | 0.25 | 2.54 | -4.73 | 5.23 | 0.924 | 1 | Wald | NM_008170.2:1788 |
| **Jag1** | 0.0106 | 0.113 | -0.211 | 0.232 | 0.927 | 1 | loglinear | NM_013822.2:2155 |
| **Bard1** | -0.0145 | 0.158 | -0.323 | 0.294 | 0.929 | 1 | lm.nb | NM_007525.3:306 |
| **Mb21d1** | -0.0211 | 0.23 | -0.472 | 0.43 | 0.93 | 1 | Wald | NM_173386.4:1068 |
| **Cd86** | 0.245 | 2.71 | -5.08 | 5.57 | 0.931 | 1 | Wald | NM_019388.3:251 |
| **Syp** | -0.0115 | 0.129 | -0.264 | 0.241 | 0.931 | 1 | lm.nb | NM_009305.2:732 |
| **Il1rap** | -0.0144 | 0.163 | -0.333 | 0.305 | 0.932 | 1 | loglinear | NM_134103.2:945 |

| | | | | | | | | |
|---|---|---|---|---|---|---|---|---|
| **Tmc7** | 0.0106 | 0.124 | -0.233 | 0.254 | 0.934 | 1 | Wald | NM_172476.4:1285 |
| **Rbfox3** | -0.00778 | 0.0917 | -0.187 | 0.172 | 0.935 | 1 | loglinear | NM_001024931.2:2700 |
| **Syn2** | -0.00849 | 0.103 | -0.211 | 0.194 | 0.937 | 1 | lm.nb | NM_013681.1:1330 |
| **Cycs** | -0.0142 | 0.183 | -0.373 | 0.345 | 0.94 | 1 | Wald | NM_007808.4:2510 |
| **Pik3cg** | 0.166 | 2.16 | -4.07 | 4.4 | 0.941 | 1 | Wald | NM_020272.2:2890 |
| **Epcam** | -0.169 | 2.22 | -4.52 | 4.18 | 0.941 | 1 | Wald | NM_008532.2:1550 |
| **Tmem206** | -0.00353 | 0.0474 | -0.0965 | 0.0894 | 0.942 | 1 | loglinear | NM_025864.3:845 |
| **Nptx1** | 0.00692 | 0.0932 | -0.176 | 0.19 | 0.943 | 1 | loglinear | NM_008730.2:4748 |
| **Entpd2** | -0.243 | 3.3 | -6.71 | 6.22 | 0.943 | 1 | Wald | NM_009849.2:1016 |
| **Tyrobp** | -0.0119 | 0.164 | -0.333 | 0.309 | 0.944 | 1 | loglinear | NM_011662.2:130 |
| **Rac2** | -0.263 | 3.75 | -7.61 | 7.09 | 0.946 | 1 | Wald | NM_009008.3:2258 |
| **Olfml3** | -0.418 | 6.08 | -12.3 | 11.5 | 0.947 | 1 | Wald | NM_133859.2:1035 |
| **H2-T23** | 0.176 | 2.56 | -4.83 | 5.19 | 0.947 | 1 | Wald | NM_010398.3:365 |
| **Kat2b** | 0.00404 | 0.0591 | -0.112 | 0.12 | 0.947 | 1 | loglinear | NM_020005.3:3030 |
| **Suv39h1** | -0.00325 | 0.0476 | -0.0966 | 0.0901 | 0.947 | 1 | loglinear | NM_011514.2:396 |
| **Ptger4** | 0.216 | 3.21 | -6.07 | 6.5 | 0.948 | 1 | Wald | NM_008965.1:315 |
| **Erbb3** | -0.246 | 3.71 | -7.51 | 7.02 | 0.949 | 1 | Wald | NM_010153.1:1290 |
| **Tnfrsf10b** | 0.343 | 5.2 | -9.85 | 10.5 | 0.949 | 1 | Wald | NM_020275.3:1625 |
| **Cd36** | 0.292 | 4.58 | -8.68 | 9.27 | 0.951 | 1 | Wald | NM_007643.3:1520 |
| **P2ry12** | 0.0117 | 0.183 | -0.347 | 0.371 | 0.951 | 1 | Wald | NM_027571.3:439 |
| **Smarca5** | 0.00349 | 0.0567 | -0.108 | 0.115 | 0.952 | 1 | loglinear | NM_053124.2:2934 |
| **Adamts16** | 0.22 | 3.68 | -6.99 | 7.43 | 0.954 | 1 | Wald | NM_172053.2:1918 |
| **Irak2** | -0.473 | 8.08 | -16.3 | 15.4 | 0.955 | 1 | Wald | NM_001113553.1:485 |
| **Pmp22** | -0.495 | 8.81 | -17.8 | 16.8 | 0.957 | 1 | Wald | NM_008885.2:395 |
| **Ccr2** | -0.152 | 2.74 | -5.52 | 5.21 | 0.957 | 1 | Wald | NM_009915.2:2965 |
| **Tmem37** | -0.353 | 6.74 | -13.6 | 12.9 | 0.96 | 1 | Wald | NM_019432.2:445 |
| **Tradd** | -0.168 | 3.32 | -6.68 | 6.34 | 0.961 | 1 | Wald | NM_001033161.2:562 |
| **Brca1** | -0.452 | 9.07 | -18.2 | 17.3 | 0.962 | 1 | Wald | NM_009764.3:2027 |
| **Hspb1** | -0.216 | 4.36 | -8.76 | 8.33 | 0.962 | 1 | Wald | NM_013560.2:630 |
| **Syk** | 0.00778 | 0.157 | -0.301 | 0.316 | 0.962 | 1 | Wald | NM_001198977.1:2064 |
| **Relb** | -0.281 | 5.72 | -11.5 | 10.9 | 0.962 | 1 | Wald | NM_009046.2:2013 |
| **Rhoa** | 0.00199 | 0.043 | -0.0822 | 0.0862 | 0.964 | 1 | loglinear | NM_016802.4:1885 |

| | | | | | | | | |
|---|---|---|---|---|---|---|---|---|
| **C5ar1** | 0.119 | 2.63 | -5.04 | 5.28 | 0.965 | 1 | Wald | NM_007577.3:595 |
| **Tgm2** | -0.326 | 7.92 | -15.9 | 15.2 | 0.968 | 1 | Wald | NM_009373.3:1260 |
| **Casp6** | -0.00573 | 0.141 | -0.283 | 0.271 | 0.969 | 1 | loglinear | NM_009811.3:360 |
| **Fbln5** | -0.169 | 4.22 | -8.44 | 8.11 | 0.969 | 1 | Wald | NM_011812.4:2138 |
| **Mavs** | 0.00515 | 0.129 | -0.247 | 0.257 | 0.969 | 1 | Wald | NM_144888.2:1162 |
| **Casp8** | -0.169 | 4.23 | -8.46 | 8.12 | 0.969 | 1 | Wald | NM_009812.2:1463 |
| **Cx3cl1** | 0.18 | 4.5 | -8.63 | 8.99 | 0.969 | 1 | Wald | NM_009142.3:125 |
| **Bola2** | -0.00132 | 0.0333 | -0.0666 | 0.064 | 0.969 | 1 | loglinear | NM_175103.3:94 |
| **Tlr4** | 0.25 | 6.31 | -12.1 | 12.6 | 0.969 | 1 | Wald | NM_021297.2:2510 |
| **Fos** | 0.0839 | 2.14 | -4.11 | 4.28 | 0.97 | 1 | Wald | NM_010234.2:1330 |
| **Csf2rb** | -0.122 | 3.19 | -6.38 | 6.14 | 0.97 | 1 | Wald | NM_007780.4:4185 |
| **Tubb3** | -0.00077 | 0.0203 | -0.0406 | 0.039 | 0.971 | 1 | loglinear | NM_023279.2:179 |
| **Ash2l** | -0.00172 | 0.0461 | -0.0921 | 0.0886 | 0.971 | 1 | loglinear | NM_001080793.1:2125 |
| **Ccl3** | -0.169 | 4.55 | -9.09 | 8.75 | 0.971 | 1 | Wald | NM_011337.1:60 |
| **Trim47** | 0.00679 | 0.185 | -0.355 | 0.369 | 0.972 | 1 | lm.nb | NM_001205081.1:2019 |
| **Dnmt3b** | -0.308 | 8.65 | -17.3 | 16.7 | 0.973 | 1 | Wald | NM_001003960.3:2312 |
| **Il2rg** | -0.226 | 6.43 | -12.8 | 12.4 | 0.973 | 1 | Wald | NM_013563.3:566 |
| **Fkbp5** | 0.00373 | 0.112 | -0.216 | 0.223 | 0.974 | 1 | loglinear | NM_010220.3:2125 |
| **Vav1** | 0.285 | 8.56 | -16.5 | 17.1 | 0.974 | 1 | Wald | NM_011691.4:1640 |
| **Ripk1** | -0.281 | 8.69 | -17.3 | 16.8 | 0.975 | 1 | Wald | NM_009068.3:1246 |
| **Kcnd1** | -0.116 | 3.6 | -7.17 | 6.94 | 0.975 | 1 | Wald | NM_008423.1:1400 |
| **Nqo1** | -0.00485 | 0.153 | -0.304 | 0.294 | 0.976 | 1 | Wald | NM_008706.5:430 |
| **Pros1** | 0.183 | 5.81 | -11.2 | 11.6 | 0.976 | 1 | Wald | NM_011173.2:2720 |
| **Cntnap2** | -0.00136 | 0.0441 | -0.0877 | 0.085 | 0.976 | 1 | loglinear | NM_001004357.2:3985 |
| **Fcrls** | -0.278 | 9.24 | -18.4 | 17.8 | 0.977 | 1 | Wald | NM_030707.3:925 |
| **Fcer1g** | -0.254 | 8.75 | -17.4 | 16.9 | 0.978 | 1 | Wald | NM_010185.4:264 |
| **Hmox1** | -0.236 | 8.14 | -16.2 | 15.7 | 0.978 | 1 | Wald | NM_010442.2:610 |
| **Fgl2** | -0.217 | 7.82 | -15.5 | 15.1 | 0.979 | 1 | Wald | NM_008013.2:3470 |
| **Bcas1** | 0.12 | 4.47 | -8.64 | 8.88 | 0.979 | 1 | Wald | NM_029815.2:932 |
| **Cd40** | -0.11 | 4.36 | -8.66 | 8.44 | 0.981 | 1 | Wald | NM_011611.2:1425 |
| **Tnfrsf1b** | 0.137 | 5.44 | -10.5 | 10.8 | 0.981 | 1 | Wald | NM_011610.3:3270 |
| **C3ar1** | 0.111 | 4.45 | -8.61 | 8.83 | 0.981 | 1 | Wald | NM_009779.2:555 |

| | | | | | | | | |
|---|---|---|---|---|---|---|---|---|
| **Il1r1** | -0.154 | 6.34 | -12.6 | 12.3 | 0.981 | 1 | Wald | NM_001123382.1:820 |
| **Pnoc** | 0.0973 | 4 | -7.75 | 7.94 | 0.981 | 1 | Wald | NM_001205075.1:332 |
| **Myct1** | -0.112 | 4.64 | -9.2 | 8.98 | 0.981 | 1 | Wald | NM_026793.2:180 |
| **Mapk12** | -0.214 | 8.85 | -17.6 | 17.1 | 0.981 | 1 | Wald | NM_013871.3:1586 |
| **Fdxr** | 0.0612 | 2.61 | -5.05 | 5.17 | 0.982 | 1 | Wald | NM_007997.1:1584 |
| **Nod1** | -0.1 | 4.36 | -8.64 | 8.44 | 0.982 | 1 | Wald | NM_172729.2:1446 |
| **Cidea** | -0.0413 | 1.82 | -3.61 | 3.52 | 0.983 | 1 | Wald | NM_007702.2:514 |
| **Prkcq** | 0.136 | 6.31 | -12.2 | 12.5 | 0.983 | 1 | Wald | NM_008859.2:1210 |
| **Bag3** | 0.00374 | 0.175 | -0.34 | 0.347 | 0.984 | 1 | Wald | NM_013863.4:1000 |
| **Bcl2a1a** | 0.0387 | 1.93 | -3.75 | 3.83 | 0.985 | 1 | Wald | NM_009742.3:175 |
| **Gal3st1** | -0.064 | 3.46 | -6.84 | 6.71 | 0.986 | 1 | Wald | NM_001177691.1:1197 |
| **Rsad2** | -0.145 | 7.85 | -15.5 | 15.2 | 0.986 | 1 | Wald | NM_021384.2:3185 |
| **Ifih1** | -0.0823 | 4.52 | -8.95 | 8.79 | 0.986 | 1 | Wald | NM_027835.2:1997 |
| **Anxa1** | 0.0687 | 3.8 | -7.37 | 7.51 | 0.986 | 1 | Wald | NM_010730.2:400 |
| **Hdac6** | 0.000891 | 0.0518 | -0.101 | 0.103 | 0.987 | 1 | loglinear | NM_010413.3:564 |
| **Cryba4** | 0.0583 | 3.47 | -6.74 | 6.86 | 0.987 | 1 | Wald | NM_021351.1:440 |
| **Egr1** | -0.071 | 4.27 | -8.44 | 8.3 | 0.987 | 1 | Wald | NM_007913.5:515 |
| **Mgmt** | 0.0807 | 5.37 | -10.5 | 10.6 | 0.988 | 1 | Wald | NM_008598.2:350 |
| **Kdm3b** | -0.0581 | 3.95 | -7.81 | 7.69 | 0.989 | 1 | Wald | NM_001081256.1:6815 |
| **Tmem119** | -0.0807 | 5.59 | -11 | 10.9 | 0.989 | 1 | Wald | NM_146162.2:1550 |
| **Fcgr3** | -0.0827 | 5.96 | -11.8 | 11.6 | 0.989 | 1 | Wald | NM_010188.5:1175 |
| **Sox10** | -0.0862 | 6.24 | -12.3 | 12.1 | 0.989 | 1 | Wald | XM_128139.6:2646 |
| **Ralb** | -0.00082 | 0.0601 | -0.119 | 0.117 | 0.989 | 1 | loglinear | NM_022327.5:1120 |
| **Gpr34** | 0.117 | 8.62 | -16.8 | 17 | 0.99 | 1 | Wald | NM_011823.4:256 |
| **Exo1** | -0.096 | 7.21 | -14.2 | 14 | 0.99 | 1 | Wald | NM_012012.4:2214 |
| **Tmem144** | -0.0596 | 4.88 | -9.62 | 9.5 | 0.991 | 1 | Wald | NM_027495.4:1525 |
| **Ddb2** | 0.0635 | 5.26 | -10.2 | 10.4 | 0.991 | 1 | Wald | NM_028119.5:94 |
| **Il6ra** | -0.0434 | 3.74 | -7.37 | 7.28 | 0.991 | 1 | Wald | NM_010559.2:2825 |
| **Ptprc** | -0.0246 | 2.81 | -5.53 | 5.48 | 0.993 | 1 | Wald | NM_011210.3:2320 |
| **Bcl2l11** | 0.0484 | 5.92 | -11.6 | 11.7 | 0.994 | 1 | Wald | NM_001284410.1:236 |
| **Tmem204** | -0.0497 | 6.22 | -12.2 | 12.1 | 0.994 | 1 | Wald | NM_001001183.1:1006 |
| **Irak4** | -0.0655 | 8.37 | -16.5 | 16.3 | 0.994 | 1 | Wald | NM_029926.5:250 |

| | | | | | | | | |
|---|---|---|---|---|---|---|---|---|
| **Tnfsf13b** | 0.0315 | 4.55 | -8.88 | 8.95 | 0.995 | 1 | Wald | NM_033622.1:225 |
| **Igf1** | 0.0268 | 3.96 | -7.73 | 7.79 | 0.995 | 1 | Wald | NM_001111274.1:418 |
| **Mpg** | -0.0454 | 6.84 | -13.5 | 13.4 | 0.995 | 1 | Wald | NM_010822.3:276 |
| **Ppp3r2** | 0.0528 | 8.41 | -16.4 | 16.5 | 0.995 | 1 | Wald | NM_001004025.4:1500 |
| **Bdnf** | -0.0458 | 7.47 | -14.7 | 14.6 | 0.995 | 1 | Wald | NM_007540.4:3260 |
| **Tle3** | -0.00038 | 0.0698 | -0.137 | 0.136 | 0.996 | 1 | loglinear | NM_009389.2:3584 |
| **Nfe2l2** | -0.0221 | 4.15 | -8.16 | 8.11 | 0.996 | 1 | Wald | NR_132727.1:144 |
| **Myd88** | 0.028 | 6.29 | -12.3 | 12.4 | 0.997 | 1 | Wald | NM_010851.2:1595 |
| **Lmna** | 0.000582 | 0.136 | -0.265 | 0.266 | 0.997 | 1 | loglinear | NM_001002011.2:1611 |
| **Slc2a5** | 0.0149 | 3.52 | -6.88 | 6.91 | 0.997 | 1 | Wald | NM_019741.3:2305 |
| **Tnfrsf12a** | 0.0198 | 4.75 | -9.29 | 9.33 | 0.997 | 1 | Wald | NM_001161746.1:517 |
| **Cd33** | -0.0185 | 5.12 | -10.1 | 10 | 0.997 | 1 | Wald | NM_001111058.1:598 |
| **Nostrin** | -0.0258 | 7.15 | -14 | 14 | 0.997 | 1 | Wald | NM_181547.3:1452 |
| **Fen1** | 0.000296 | 0.0853 | -0.167 | 0.167 | 0.997 | 1 | loglinear | NM_001271614.1:1880 |
| **Kdm5b** | -7.28E-05 | 0.0441 | -0.0866 | 0.0865 | 0.999 | 1 | loglinear | NM_152895.2:3620 |
| **Man2b1** | -0.00013 | 0.0983 | -0.193 | 0.193 | 0.999 | 1 | loglinear | NM_010764.2:1658 |
| **Plcg2** | -0.00319 | 6.62 | -13 | 13 | 1 | 1 | Wald | NM_172285.1:978 |
| **Socs3** | 0.00238 | 8.67 | -17 | 17 | 1 | 1 | Wald | NM_007707.2:585 |
| **Arhgap24** | -1.33E-15 | 0.154 | -0.301 | 0.301 | 1 | 1 | lm.nb | NM_029270.2:1164 |

Differential expression (IAV vs. Con, Log2 fold change) of 591 genes in fetal brain tissue of litters from saline control and IAV-infected gestating dams at GD17 (1 fetal brain/litter; Con: n = 6, IAV: n = 4). Raw Data were analyzed using the Advanced Analysis feature of the nSolver 4.0 software, with automated normalization gene selection and quality control measures (27 housekeeping and positive and negative control probes included; 166 probes excluded based on the “Omit Low Count Data” default selection).
