## Supplementary material for "Moderately pathogenic maternal influenza A virus infection disrupts placental integrity but spares the fetal brain": MIA reporting guidelines checklist

**Maternal Immune Activation Model Reporting Guidelines Checklist**

| ARRIVE Reporting Guideline & Recommendation | Arrive Item | MIA Model Specific Reporting Recommendation<br>***Please complete this chart for each point outlined below. If not applicable, write N/A*** |
|---|---|---|
| **Study design**<br>➢Overview of immune activation issues<br><br>For each experiment, give brief details of the study design including:<br>a. The number of experimental and control groups.<br>b. Any steps taken to minimize the effects of subjective bias when allocating animals to treatment (e.g. randomization procedure) and when assessing results (e.g. if done, describe who was blinded and when).<br>c. The experimental unit (e.g. a single animal, group or cage of animals).<br><br>A time-line diagram or flow chart can be useful to illustrate how complex study designs were carried out. | 6 | MIA Specific Reporting:<br>a. General need for improved reporting in MIA model methods + reporting pilot data<br>○ Details on pilot data:<br><br>While we have previously validated the same strain and dose of mouse-adapted IAV (H3N2) in non-pregnant male mice, infection of pregnant and non-pregnant female mice was piloted in two separate cohorts of mice (using 1000 TCID50 of X31).<br>Cohort 1: Twelve nulliparous adult C57BL/6 female mice were pair-housed and bred as described in the Methods. Non-pregnant and confirmed pregnant females were both inoculated with IAV X31 as described, with two non-pregnant females serving as controls. Sickness symptoms and weight were monitored daily, and all animals were sacrificed 8 dpi and tissues collected. Circulating immune cells, lung and reproductive tissues were examined to determine infection severity and confirm fetal viability.<br>Cohort 2: Eight nulliparous adult C57BL/6 female mice were pair-housed, bred, confirmed pregnant, then randomly assigned to Con and IAV groups (n = 4/group) and inoculated as described in the Methods. Sickness symptoms and weight were monitored daily, and all dams were allowed to gestate through parturition to confirm normal gestation length, pup viability, and maternal rearing ability. Offspring were aged out to adulthood. |
| **Experimental procedures**<br>➢Compounds<br>➢Validation measures<br><br>For each experiment and each experimental group, including controls, provide precise details of all procedures carried out. For example:<br>a. How (e.g. drug formulation and dose, site and route of administration, anaesthesia and analgesia used [including monitoring], surgical procedure, method of euthanasia). Provide details of any specialist equipment used, including supplier(s).<br>b. When (e.g. time of day).<br>c. Where (e.g. home cage, laboratory, water maze).<br>d. Why (e.g. rationale for choice of specific anaesthetic, route of administration, drug dose used). | 7 | Provide details of:<br>a. Compounds – source, vehicle, preparation/storage, administration route, volume administered, whether anesthetics were used at time of immune challenge.<br>○ Name of compound: Mouse-adapted influenza A virus H3N2 X31<br>○ Catalogue number: N/A<br>○ Lot number: N/A<br>○ Vehicle control used: sterile D-PBS<br>○ Route of administration: Intranasal<br>○ Volume administered: 1000 TCID50 in 50 µL<br>○ Storage conditions: Viral stock kept at -80°C<br>○ Anesthetic (type, dose, duration) used:<br>Inhalation of isoflurane (gas, 2.5%, 30-45 seconds)<br>b. Housing variables at injection - temperature of room at injection time, cage change at time of injection or not<br>○ Light cycle of animal housing room: 12-hour, 06:00 - 18:00 light<br>○ Time of day of injection: morning<br>○ Room temperature at injection time: ambient (22°C)<br>○ Did a cage change occur at time of injection: NO |

Kentner AC, Bilbo AD, Brown AS, Hsiao EY, McAllister AK, Meyer U, Pearce BD, Pletnikov MV, Yolken RH, Bauman MD. (2018). Maternal immune activation: reporting guidelines to improve the rigor, reproducibility, and transparency of the model. Neuropsychopharmacology, https://doi.org/10.1038/s41386-018-0185-7.

| | | |
|---|---|---|
| | | c. Validation of immune activation – behavior, physiological indices and/or cytokine data, including pilot dosing data<br>○ Method used to verify immune activation:<br>Immune activation was verified and previously reported in male mice (Kenney et al. 2019, PNAS, doi:10.1073/pnas.1900784116). Pilot data on IAV infection in female mice (pregnant and non-pregnant) was validated through sickness behaviors, failure to gain weight, changes in circulating immune cell populations (flow cytometry), and lung inflammation (qPCR).<br>d. Validation of gestational timing – vaginal plug, estrous cycle, weight gain<br>○ Method of validating gestational timing:<br>Estrous cycle was initiated by exposing virgin females to soiled male bedding, the male was introduced to the females on the third day, and vaginal plug was used to determine successful copulation and was designated GD1. Weight gain of 2 grams or more by GD10 was used to confirm successful pregnancy.<br>Additional comments:<br>Presence of a vaginal plug was assessed each morning (~07:00) for four consecutive days after introduction of the male stud, then females were moved back to a clean cage. |
| **Experimental animals**<br>➢Species/strain/vendor<br>a. Provide details of the animals used, including species, strain, sex, developmental stage (e.g. mean or median age plus age range) and weight (e.g. mean or median weight plus weight range).<br>b. Provide further relevant information such as the source of animals, international strain nomenclature, genetic modification status (e.g. knock-out or transgenic), genotype, health/immune status, drug or test naïve, previous procedures, etc. | 8 | Provide details of:<br>a. Species – considerations for appropriate species (mouse, rat, non human primate, other)<br>○ Species: Mouse<br>b. Strain – variability in strain can influence model<br>○ Strain: C57BL/6NTac<br>c. Maternal/Offspring Physiological Variables at time of immune challenge – age, body weight<br>○ Maternal Age at challenge: 12 - 38 weeks<br>○ Maternal Body weight: 19-28 grams<br>○ Offspring Age at challenge: No secondary challenge<br>○ Offspring Sex: Both males and females tested<br>○ Offspring Body weight: N/A<br>d. Vendor – even within the same strain, vendor can influence endpoints<br>○ Vendor: Taconic Biosciences<br>○ Location of Vendor: Cambridge City, IN, USA<br>○ Room/area where animals originated from: N/A |

| | | |
|---|---|---|
| | | Additional Comments:<br>Maternal age at challenge was variable because of circumstances that were out of our control due to the COVID19 pandemic, which shut down our research effort for 3 months. It was our intention that all dams be challenged at 12 - 15 wks old, but we were forced to extend this age limit. |
| **Housing and husbandry**<br>➢Cage, ventilation, bedding, enrichment<br><br>Provide details of:<br>a. Housing (type of facility e.g. specific pathogen free [SPF]; type of cage or housing; bedding material; number of cage companions; tank shape and material etc. for fish).<br>b. Husbandry conditions (e.g. breeding program, light/dark cycle, temperature, quality of water etc for fish, type of food, access to food and water, environmental enrichment).<br>c. Welfare-related assessments and interventions that were carried out prior to, during, or after the experiment. | 9 | Provide details of:<br>a. Caging systems<br>○ *At breeding*<br>Material of cage: individually ventilated microisolator polysulfone<br>Cage dimensions: 15.01" L x 5.13" W x 7.63" H<br>○ *After parturition*<br>Material of cage: N/A<br>Cage dimensions: N/A<br>○ *At weaning*<br>Material of cage: N/A<br>Cage dimensions: N/A<br><br>b. Animal Holding room<br>○ Temperature in room: ambient (22°C)<br>○ Humidity in room: 45%<br>○ Ventilation system: Positive pressure<br>○ Specific pathogen free [SPF]: YES<br>○ Are males & females housed in the same or separate rooms: Housed in same room after weaning<br>c. Bedding exchanges/bedding type<br>○ Type of cage bedding used: corn cob (Bed-o'Cobs brand)<br>○ Frequency of cage changes per week<br>*during gestation:* 2 cage changes total during gestation<br>*during neonatal period:* N/A<br>*following weaning:* N/A<br><br>d. Breeding - bred on site or timed pregnant, how many different sires (are the same fathers breeding with both experimental and control dams)<br>Breeding location: bred on site, 15 different sires bred to Con and IAV dams |

- Gestational age at shipping: N/A
- Biological age of dams (if not listed in Section 8c): see 8c
- Number of Dams bred: 71
- How many times have dams been mated previously: 0
- How many times did the dams mate and not become pregnant: 0-3
- Are the dams primiparous or multiparous? All dams are primiparous
- What was the frequency of maternal handling during the gestational/neonatal period (e.g. cage cleanings, weighing, blood collection manipulations): daily handling for weighing only
- Biological age of sires: 12 - 38 wks
- Number of sires bred: 15
- How many times have sires been mated previously: 0 to 3
- How many times did the sires mate successfully (e.g. mating resulted in pregnancy, full term birth): 1 to 4
- If bred previously, what was the interval between mating times: >4wk
- Are sires matched to experimental and control dams: YES
- Describe the mating design (1:1, 1:2 etc): 1:2

e. Social enrichment – number of cage companions
   - Number of cage companions prior to breeding: pair housed
   - Gestational age when dam separated for parturition: GD 10
   - Number of cage companions at weaning: N/A

f. Physical enrichment – describe enrichment devices, and when enrichment is in the cage (removed when pups born? Or present throughout study), does the enrichment type change? How frequently?
   - Describe what type of enrichment devices (and how many) are included in cage/housing room:

Pair-housed females were given a cotton nestlet with which to assemble a nest. If barbering occurred, crinkled paper was added. The nest material was present prior to and post-breeding, until GD10, when females were singly housed.

Singly-housed pregnant females were given a plastic hut and cotton nestlet. This hut and nest material was present throughout the study (GD10-17).

All cages had standard bedding and nest material throughout the study.

| | | |
|---|---|---|
| | | ○ Does enrichment type/access change across study? YES<br>○ If so, when does enrichment type/access change (e.g. enrichment removed prior to parturition and replaced in late neonatal period):<br>As described above: when pair-housed, there is no "additional" enrichment outside nest material and companionship. Once singly-housed (GD10-GD17), a plastic enrichment hut is added.<br><br>Additional Comments: |
| **Sample size**<br>➢Litter versus offspring<br><br>a. Specify the total number of animals used in each experiment, and the number of animals in each experimental group.<br>b. Explain how the number of animals was arrived at. Provide details of any sample size calculation used.<br>c. Indicate the number of independent replications of each experiment, if relevant. | 10 | Provide details of:<br>a. Maternal N vs offspring N<br>○ What is the total number of dams/litters included in the study: 37<br>○ What is the total number of offspring per litter included the study: 4 fetuses per litter, 4 placentas per litter<br>b. Litter size and sex distribution<br>○ What size was each litter maintained at: N/A<br>○ What age did culling take place at: N/A<br>○ How many males and females were maintained in each litter: N/A<br>c. Cross fostering<br>○ Did cross fostering occur: NO<br>○ If so, at what age did cross fostering occur: N/A<br><br>Additional Comments: |

| | | |
|---|---|---|
| **Allocating animals to experimental groups**<br><br>a. Give full details of how animals were allocated to experimental groups, including randomization or matching if done.<br>b. Describe the order in which the animals in the different experimental groups were treated and assessed. | 11 | a. How many offspring per litter were used in each measure: 1-2<br><br>b. Randomization/Matching procedures<br>○ What procedures were used to assign animals to groups:<br>Fetuses were randomly assigned numbers (starting with 1) during tissue collection, and then tissue was assigned to different outcomes going in order (ex: fetal brain #1 from each litter was used for nanoString gene expression, fetal brain #2 was used for qPCR, #3 for Proteome Profiler; placentas #1 and #2 from each litter were used for histology, placenta #3 for Proteome Profiler, etc.)<br><br>c. Sex as a biological variable (behavioral and physiological outcomes)<br>○ Were both males and females evaluated in each behavioral and physiological outcome: YES<br><br>Additional Comments:<br>Fetal sex was not identified; however, because fetuses were randomly assigned numbers at collection, and because 4 representative fetuses per litter were used, we can assume both male and female fetuses were included in each measure. |
| **Experimental outcomes**<br>➢ Behavioral testing<br>➢ Physiological endpoints<br><br>Clearly define the primary and secondary experimental outcomes assessed (e.g. cell death, molecular markers, behavioral changes). | 12 | a. Maternal behavior and pup interactions<br>○ If maternal care was evaluated, were there differences following immunogen challenge (if so, please briefly describe):<br>N/A<br><br>b. Age(s) of offspring at behavioral testing/physiological evaluation endpoints:<br>N/A<br><br>c. Order of testing (e.g. behavioral test order)<br>○ Were animals evaluated in a counter-balanced order in terms of:<br>*presentation of tests to each animal:*<br>*order of experimental/control groups run through each test:*<br>○ What was the inter-test interval if a single animal underwent a battery of tests:<br>N/A |

| | | |
|---|---|---|
| | | Additional Comments: |
| **Statistical methods**<br><br>a. Provide details of the statistical methods used for each analysis.<br>b. Specify the unit of analysis for each dataset (e.g. single animal, group of animals, single neuron).<br>c. Describe any methods used to assess whether the data met the assumptions of the statistical approach. | 13 | a. Unit of analysis for each data set<br>○ Is the unit (n) of each analysis based on number of litters, or number of animals used per group:<br><br>Number of litters/dams |
| **Other Disclosures** | | Please make note of any other extraneous variables that you would like to report (e.g. fire alarms, construction, temporary relocations, other variables that you think we should be considering in our studies etc.):<br><br>Unforeseen limitations arose during experimentation due to the COVID19 pandemic. However, no replicate was directly impacted or interrupted by the shut down. Rather, the time between the first two replicates and the last two replicates was delayed 3 months (thus, each replicate still began and was completed under identical experimental conditions). |

The recommended use of this reporting form is to fill it out and include it as supplemental material for each of your laboratory's research publications. If there are difficulties utilizing/adapting this fillable form, please contact one of the corresponding authors to request a copy. The authors give permission for this table to be edited for use in reporting on other animal models (e.g. postnatal immune challenge models, early life stress models) as appropriate.

Kentner AC, Bilbo AD, Brown AS, Hsiao EY, McAllister AK, Meyer U, Pearce BD, Pletnikov MV, Yolken RH, Bauman MD. (2018). Maternal immune activation: reporting guidelines to improve the rigor, reproducibility, and transparency of the model. Neuropsychopharmacology, https://doi.org/10.1038/s41386-018-0185-7.
